# Controlling gene expression in mammalian cells using multiplexed conditional guide RNAs for Cas12a

**DOI:** 10.1101/2021.04.16.440136

**Authors:** Lukas Oesinghaus, Friedrich C. Simmel

## Abstract

Spatiotemporal control of the activity of Cas proteins is of considerable interest for both basic research and therapeutics. Only few mechanisms have been demonstrated for regulating the activity of guide RNAs (gRNAs) for Cas12a in mammalian cells, however, and combining and compactly integrating multiple control instances on single transcripts has not been possible so far. Here, we show that conditional processing of the 3’ tail is a viable general approach towards switchable Pol II-transcribed Cas12a gRNAs that can activate gene expression in mammalian cells in an input-dependent manner. Processing of the 3’ tail can be achieved using microRNA and short hairpin RNA as inputs, via a guanine-responsive ribozyme, and also using an RNA strand displacement mechanism. We further show that Cas12a along with several independently switchable gRNAs can be integrated on a single transcript using stabilizing RNA triplexes, providing a route towards compact Cas12a-based gene regulation constructs with multi-input switching capabilities.

## Introduction

The CRISPR-associated proteins Cas9 and Cas12a are RNA-guided DNA nucleases^1,2^, while their DNase-dead variants (dCas9 and dCas12a) are programmable DNA binders that can be fused to additional domains for transcriptional activation and repression, base editing, and other functions in mammalian cells^3–5^. Both for basic research and therapeutic purposes, it is desirable to exert spatiotemporal control on their activity^6^. This can be achieved via inducible or tissue-specific promoters, additional components such as anti-CRISPR proteins, or through switchable CRISPR components whose activity is contingent on control inputs such as light, small molecules or endogenous RNAs^6^. For switchable control mechanisms, there are three fundamental options: either the Cas protein itself can be made switchable, its associated mRNA, or its guide RNA (gRNA). For interfacing with endogenous RNAs such as miRNAs or other mRNAs, messenger and guide RNAs are natural choices as they can be addressed via predictable Watson-Crick base pairing. Using gRNA as a switchable element has the additional advantage that a combination of multiple different targets and inputs in a single experiment can be achieved comparatively easily. Using Cas protein orthologs, for example, requires delivering a separate protein for each individual target^7^. Delivering multiple gRNAs is much simpler and more compact, i.e., it requires far less total sequence length. Cas12a, specifically, processes its own gRNA arrays, which makes it straightforward to place multiple gRNAs on a single transcript^8^. Using a triplex-forming sequence from the Malat1 noncoding RNA to stabilize the protein coding sequence, Campa et al. recently managed to place Cas12a along with 15 different gRNAs on a single Pol II transcript^9^.

Over the past years, multiple strategies have been developed to engineer switchable gRNAs that respond to a wide range of exogenous and endogenous triggers. Cas9 gRNAs have been switched with small molecules using aptamers embedded in the gRNA structure^10,11^ or using inducible self-cleaving ribozymes at the 5’ end of the gRNA that interact with the spacer sequence^12^. For Cas12a, spacer-interacting riboswitches were implemented^13^. Wang et al. demonstrated microRNA-responsive Cas9 gRNAs by placing microRNA target sites adjacent to the gRNA on a Pol II transcript^14^.

Recently, several papers demonstrated activation of Cas9 or Cas12a gRNAs using toehold-mediated strand displacement^15–20^. Most of these gRNAs did not have independent input and target sequences, were not implemented in mammalian cells, or required several separate components to be delivered and expressed. Several of these shortcomings have been addressed individually. Our own earlier design of switchable Cas12a gRNAs circumvented sequence constraints using an RNA helper strand that connected an arbitrary input sequence to the switching sequence^19^. The design developed by Jin et al. implemented switchable Cas9 gRNAs without sequence constraints, but has not been shown to work in either bacteria or mammalian cells^17^. Recently, two reports have solved two of these problems simultaneously: Collins et al. developed an approach to directly sense natural transcripts using engineered Cas12a gRNAs without the use of helper strands in E. coli cells^20^, while Lin et al. demonstrated activation of Cas9 gRNAs in mammalian cells without formal sequence constraints by using two U6-transcribed helper strands^18^. So far, however, no design has been shown to address these three issues simultaneously.

Engineering multiple, independently switchable gRNAs for gene expression in mammalian cells is challenging due to the large size of the eukaryotic promoter regions, which are much larger than the gRNAs they generate. Production of several gRNAs would require several promoters on a correspondingly large plasmid, or the use of several plasmids in parallel, making true multiplexing difficult.

In the present work, we overcome this problem by combining switchable gRNAs with a design strategy that works with a single transcript for the Cas12a protein and its gRNAs. We start out by demonstrating that conditional processing of the 3’ tail of gRNA-containing CAG promoter transcripts is a general method to generate switchable Cas12a gRNAs. In particular, we show activation of gRNA activity using three different mechanisms: first, we use target sites for endogenous microRNAs and externally delivered short hairpin RNAs (shRNAs) to cleave off the 3’ tail, we then use a guanine-inducible self-cleaving ribozyme for this purpose, and finally we engineer a novel pseudoknot-based strand invasion mechanism that switches the conformation of the gRNA handle structure and thus enables cleavage by Cas12a’s own RNase activity. Thus separating the sequence region responsible for switching from the active gRNA structure avoids the sequence constraints of our earlier designs^19^. Furthermore, insulation of the Cas12a coding sequence and multiple switchable gRNA sequences through the repeated use of an RNA triplex structure enables the integration of a complete Cas12a gRNA circuit on a single transcript.

### MicroRNA- and shRNA-responsive Cas12a gRNAs

Cas12a gRNAs have previously been placed downstream of protein-coding sequences on mRNAs, where the intrinsic RNase activity of Cas12a excises them from the transcript^21^. To generate an active gRNA molecule, an additional handle sequence needs to be added downstream of the active gRNA sequence to remove the 3’ tail. As a first step towards our implementation of switchable gRNAs, we verified that trimming of a gRNA-containing transcript also works when transcribing Cas12a gRNAs from a CAG promoter without any protein-coding sequence. As a readout, we use a secreted nanoluciferase under the control of a minimal promoter containing seven repeats of a 20 nt long target sequence (t1 or t2) in HEK293 cells, which is in turn activated by an enhanced AsdCas12a-VPR (Acidaminococcus DNase-dead Cas12a-VPR) fusion (Figure 1a)^3^. The Cas12a, nanoluciferase, and gRNA are delivered on separate plasmids (Supplementary Figure 1). As expected, the gRNA is highly active when the 3’ tail is removed by Cas12a’s RNase activity, although its “OFF” activity without removal of the tail is still significantly higher than for a conventional U6-transcribed gRNA with a non-targeting spacer (Figure 1b). We reasoned that processing of the 3’ tail could be engineered to become conditional on external or endogenous molecular inputs, and thus the gRNA would become switchable by that input. As such, any conditional RNA cleavage mechanism would potentially result in switchable gRNAs.

**Figure 1.**
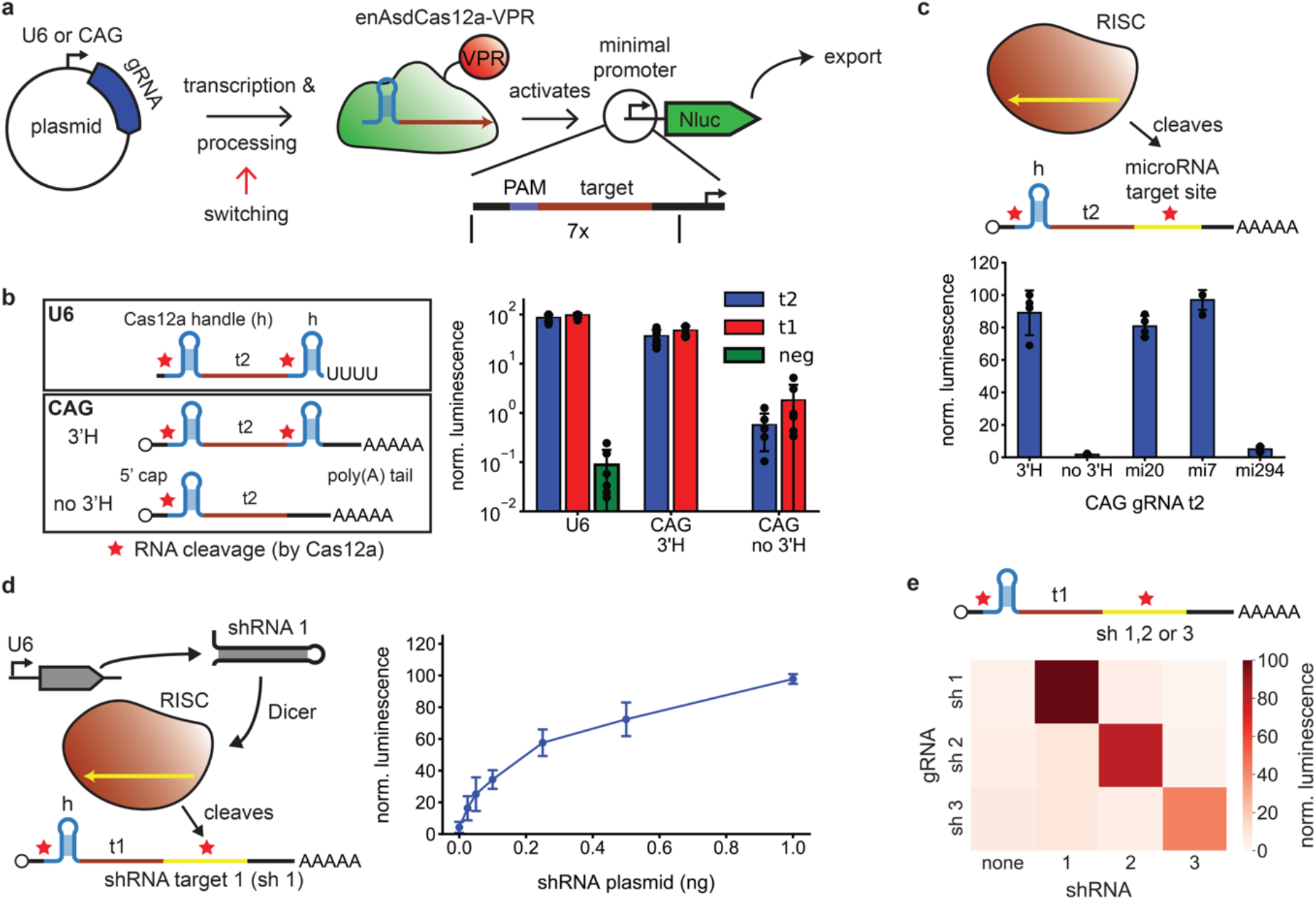
MicroRNA- and shRNA-responsive Cas12a gRNAs. **a** Guide RNAs are transcribed using a U6 or CAG promoter. The switching principle used in this study is based on conditional post-transcriptional processing of CAG promoter-transcribed gRNAs. Cas12a gRNA activity is measured via the activation of expression of secreted nanoluciferase by enAsdCas12a-VPR (enhanced Acidaminococcus DNase-dead Cas12a-VPR). **b** Nanoluciferase assay of the gRNA activity of U6-transcribed gRNAs with three different target sequences, and CAG-transcribed gRNAs with and without a 3’ gRNA handle for two different target sequences. The red star denotes an expected cleavage site, here due to Cas12a’s RNA processing activity. t2, t1 = cognate spacer sequences, neg = non-targeting spacer sequence (N=6) **c** Nanoluciferase assay of gRNAs in which the 3’ gRNA handle sequence was replaced by a microRNA target sequence. (N=4) **d** Activation of CAG gRNA t1 by a shRNA transcribed using the U6 promoter normalized to the activity of a gRNA with a 3’ handle sequence. (N=4) **e** Three gRNAs containing different shRNA targets are activated by three different shRNAs. (N=4)

A recent study by Wang et al. has demonstrated that putting two microRNA (miRNA) target sites adjacent to a Cas9 gRNA on a (CAG promoter-transcribed) Pol II-transcript results in miRNA-responsive gRNAs^14^, presumably via cleavage of the transcript by the RNA-induced silencing complex (RISC). As our first conditional processing mechanism, we therefore also attempted to control our Cas12a gRNAs by placing microRNA target sites adjacent to the target sequence (Figure 1c). As expected, hsa-miR-20a-5p and hsa-let-7a-5p target sites, whose corresponding microRNAs are expressed in HEK293 cells^22^, lead to a strong activation of gRNA activity. Conversely, a miRNA that is not expected to be present (mmu-miR-294-5p)^14^, does not. As in this case, we generally observed a low level of leaky activation for some uncleaved 3’ sequences, while others appeared to be entirely inactive in all experiments. We verified the results using flow cytometry, for which the sequence of a fluorescent protein (mScarlet) was placed downstream of the minimal promoter (Supplementary Figure 2). Qualitatively, we observe the same behavior as for the nanoluciferase assay, but with an even better on/off ratio and a lower leak for the miR-294 sequence.

Alternatively, we also tested conditional activation using a short hairpin RNA (shRNA) transcribed from a U6 promoter (Figure 1d). The shRNA was designed using software by Gu et al.^23^ to target the sequence directly downstream of the spacer sequence of gRNA t1 without a 3’ handle (cf. Figure 1b). At just 1 ng of shRNA plasmid, the gRNA activity already reaches the level achieved with the gRNA construct containing a 3’ handle sequence. We surmise that this high level of activity is likely due to multi-turnover cleavage of gRNA tails by the RISC. We designed two additional shRNAs and placed their targets downstream of two additional CAG-transcribed gRNAs with a t1 spacer sequence (Figure 1e). All gRNAs are activated by their respective shRNA, but not by the other shRNAs.

### Small molecule-responsive Cas12a gRNAs using an inducible ribozyme

As a second mechanism, we used transcript processing by ribozymes. Small molecule-inducible ribozymes have been used extensively to make switchable mammalian mRNAs^24–26^. There, activation of the ribozymes deactivates the mRNA due to degradation in the cytosol after removal of the poly(A) tail. For Pol II-transcribed gRNAs, we expect the opposite effect, namely activation of the gRNA by activation of the ribozyme.

In order to test the basic principle, we first inserted a constitutively active hepatitis delta virus (HDV) ribozyme^27^ downstream of the gRNA spacer sequence. As for the Cas12a 3’ handle construct (Figure 1b), a HDV ribozyme at the 3’ end of the gRNA indeed leads to activation, while a mutated, inactive ribozyme results in an inactive gRNA (Figure 2a). To make the gRNAs respond to small molecules, we next used a guanine-responsive HDV ribozyme originally developed by Nomura et al. for switchable mRNAs (Figure 2a)^27^. The nominally best performing ribozyme (GuaM8HDV) initially showed poor performance when inserted directly after the gRNA spacer sequence. While the activated ribozyme (at 100 μM guanine in the medium) has an activity similar to the wild-type ribozyme, there is also strong gRNA activity in the absence of guanine. In order to suppress potential interactions between the gRNA sequence and the ribozyme, we introduced a clamp structure to separate the different sequence domains and facilitate correct folding of the ribozyme (Supplementary Figure 3a and b, GuaM8 v4 in Figure 2a). With our best-performing design, an on/off ratio of 34 could be achieved, but the total activity was significantly less than for the wild-type ribozyme.

**Figure 2.**
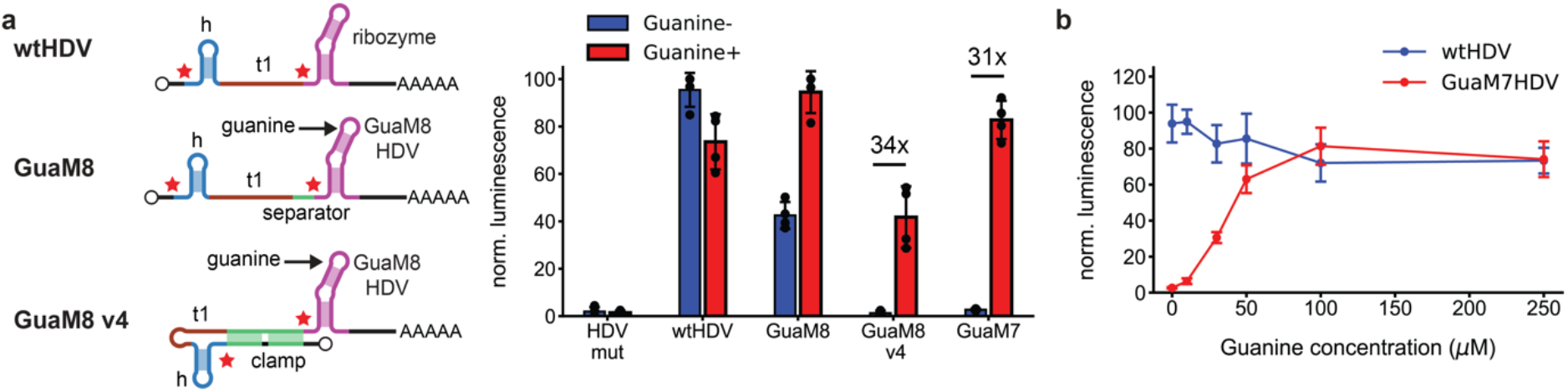
Small molecule-responsive Cas12a gRNAs using an inducible ribozyme. **a** Nanoluciferase assay of activity for CAG-transcribed gRNAs containing different wild-type or guanine-responsive ribozymes. The activity was induced with 100 μM guanine. (mut = mutated, inactive ribozyme. The red star indicates cleavage of the RNA.) (N=4) **b** Activation curve for gRNAs containing either the wtHDV or the guanine-responsive GuaM7HDV at the 3’ end. (N=4)

We also tested an alternative ribozyme (GuaM7HDV)^27^, which had a lower activity in terms of mRNA knockdown, but also less leak than GuaM8HDV. Remarkably, in the context of our conditional gRNA GuaM7HDV achieved an on/off ratio of around 31 with full total activity without any clamping or optimization, and with an activity saturating at around 100 μM guanine input (Figure 2b). As discussed in the Supporting Information, the different behaviors of the GuaM8HDV and GuaM7HDV constructs are not obvious from simulations of their secondary structures (Supplementary Figure 3c), but may be the consequence of different optimization objectives for mRNA *inactivation* and gRNA *activation*, respectively (Supplementary Note 1).

### Activation of gRNA processing using toehold-mediated strand displacement

As shown above (Figure 1), CAG promoter-transcribed gRNAs equipped with a second handle sequence at the 3’ end can be processed into a fully active form via Cas12a cleavage. We had previously demonstrated^19^ that the handle sequence itself can be sequestered into an inactive secondary structure, which can be opened by a trigger RNA via a toehold-mediated strand invasion process. This allows the handle to fold and thus enables Cas12a binding and processing of the gRNA.

We here revisited this concept in the context of the CAG promoter-transcribed gRNAs to facilitate conditional gRNA activation via 3’ handle cleavage. To avoid the sequence constraints of our earlier design, we developed a new switching principle that utilizes the fact that Cas12a recognizes the handle of its gRNA via a pseudoknot structure formed between the 5’ end of the handle and its stem-loop (Figure 3a)^28^. Accordingly, we expected disruption of the pseudoknot to be sufficient to prevent Cas12a from binding even when the handle hairpin is otherwise correctly formed. Using the nucleic acid design tool NUPACK, we generated a series of designs for pseudoknot-switchable handle structures^29^ (see also Supplementary Note 2 and Supplementary Software).

**Figure 3.**
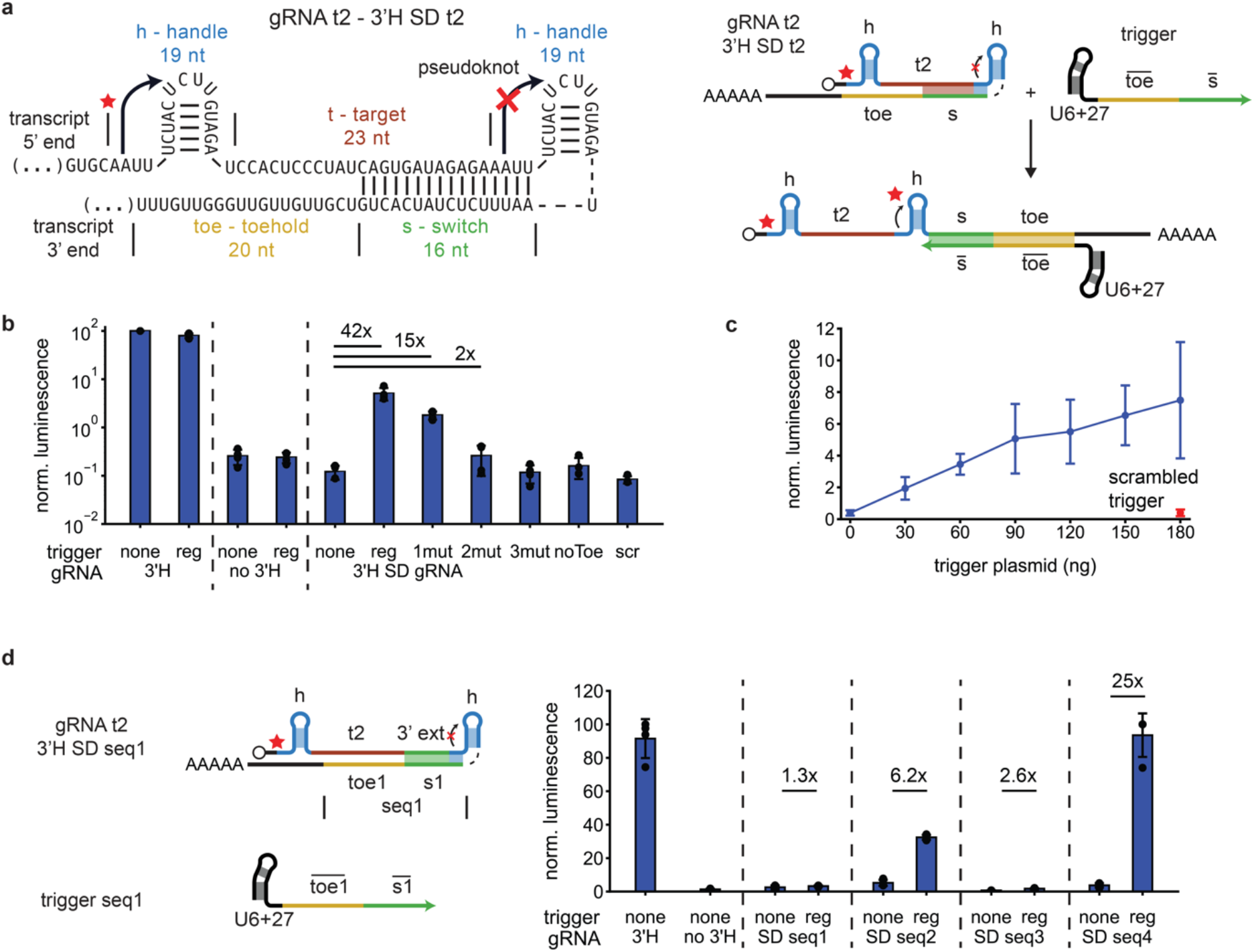
Switching of a 3’ Cas12a handle via strand displacement (3’H SD). **a** The switching principle is based on suppression of pseudoknot formation in the 3’ handle. Displacement of the secondary structure by an RNA trigger molecule allows for pseudoknot formation and therefore processing by Cas12a (red star). **b** Nanoluciferase assay of the activation of different gRNAs by U6+27-transcribed trigger molecules. (gRNAs:3’H – regular gRNA with 3’ handle, no 3’H – regular gRNA without 3’ handle, 3’H SD gRNA – gRNA with suppressed 3’ handle. triggers: reg (regular) – fully complementary trigger for the 3’H SD design, 1/2/3 mut: trigger with 1/2/3 changed nucleotides in the s domain, noToe – trigger with scrambled toehold, scr – completely scrambled trigger.) (N=4) **c** Activation of the 3’H SD gRNA shown in a with different amounts of trigger. (N=8) **d** Nanoluciferase assay of the activation of 3’H SD gRNAs with different de-novo designed trigger sequences (seq 1-4) transcribed by the U6+27 promoter. (N=4)

We first studied the basic mechanism in in vitro experiments using purified RNA, for which we paired the target sequence of the gRNA with a switch sequence located at its 5’ end to suppress pseudoknot formation (Supplementary Figure 4a). As expected, the resulting gRNA is inactive and becomes active after addition of a trigger that displaces the switch (Supplementary Figure 4b, c). As with the other switching mechanisms described above, the separation between the structure of the active gRNA and the sequence responsible for the cleavage mechanism considerably simplifies the design. As the input molecule does not need to interact directly with the sequence of the handle, no sequence constraints on the trigger molecule are incurred. This allows, in principle, to sense any accessible subsequence of a full mRNA transcript (Supplementary Figure 4b, c).

For our implementation in mammalian cells, we moved the switch and toehold domains to the 3’ end of the switched gRNA (Figure 3a), a design which we refer to as a 3’ handle SD gRNA (3’H SD gRNA). Transcription of a trigger from a U6+27 promoter^30^ alongside the 3’H SD gRNA leads to a 42-fold activation of gene expression via enAsdCas12a-VPR (Figure 3b). Changing even a single nucleotide in the switch domain of the trigger – and thus inhibiting the strand displacement process – reduces this to 15-fold activation, changing two nucleotides reduces activation to 2-fold, while three changed nucleotides reduce it to the background level. A trigger with a non-cognate toehold and a trigger with a completely scrambled sequence fail to activate the 3’H SD gRNA.

Activation of the 3’H SD gRNA by the trigger scales approximately linearly with the amount of trigger, but displays considerable variability from experiment to experiment (Figure 3c). Although the ON/OFF ratios achieved with activation by strand displacement are comparable to those obtained by the other approaches shown above, with ≈ 6-10% the maximum activation level is significantly less than that obtained with a regular control gRNA.

Another drawback of the design described in Figure 3a are its strong sequence constraints, as 12 nt of the 5’ gRNA spacer and the first four nucleotides of the 3’ handle (AAUU) are used as the switch sequence. The sequence constraints can be alleviated by removing the four handle-complementary nucleotides from the trigger (reducing activation by a factor of slightly less than two (Supplementary Figure 5a)), and by inserting an additional sequence domain between 5’ gRNA target and 3’ handle domain (cf. Supplementary Figure 6a). The resulting design retains no formal sequence constraints and therefore allows, in principle, sensing of arbitrary RNA inputs without the need for additional adaptor strands. Surprisingly, several iterations of the unconstrained design failed or did not result in appreciable ON/OFF ratios (Supplementary Note 3, Supplementary Figures 6–8), which we attributed to potential target-toehold interactions.

We therefore designed four 3’H SD gRNAs with different switch and toehold sequences, for which we explicitly excluded undesired interactions of the toehold and the full expected gRNA transcript (Supplementary Figure 9). Interestingly, the four sequences had a widely varying performance (Figure 3), ranging from non-functional (seq 1) to an excellent 25-fold activation (seq 4) by the trigger, and with a maximum activity comparable to the level of the positive control.

While the good performance of seq 4 demonstrates that it is possible to utilize our design for toehold-mediated strand displacement for CRISPR-based gene activation in mammalian cells, we were not yet able to find a clear design-performance relationship for the switches. In particular, the badly performing sequences 1 and 2 show the most stable folding, while sequence 4 has the worst predicted folding behavior (Supplementary Figure 9). At this point, our design approach does not consider the influence of the cellular environment on the folding of the gRNA switches, and we also did not account for interactions with RNA-binding proteins and with the transcriptome. It seems likely that a better understanding of these factors will be required to enable a more reliable design of mammalian 3’H SD gRNAs for arbitrary inputs and targets.

### Encoding Cas12a and multiple switchable gRNAs on a single transcript

Compared to other approaches towards multiplexed gene regulation, in which different target sites are addressed with different stimuli, switchable gRNAs have the potentially major advantage that they do not necessarily require the delivery of multiple large gene constructs. In the examples described so far, however, only a single conditional gRNA was delivered on a separate plasmid than the dCas12a gene. We therefore sought to develop a strategy that allows the integration of the dCas12a coding sequence together with multiple switchable gRNAs on a single transcript.

While it is possible to simply place several regular (non-switchable) Cas12a gRNAs next to each other^9,31^, this straightforward approach fails for our switchable gRNAs (Figure 4a). The gRNA located at the 3’ end of the transcript works as desired, retaining a good ON/OFF ratio and total activity. However, the gRNA at the 5’ end has a drastically reduced activity in the ON state. We surmise that the removal of the poly(A) tail during transcript processing leads to an enhanced degradation of the RNA, which affects the function of the 5’ gRNA domain more than that of the 3’ gRNA. This interpretation is in agreement with a set of experiments, in which we systematically varied the 3’ sequence context (Supplementary Figure 10 and 11, Supplementary Note 4).

**Figure 4.**
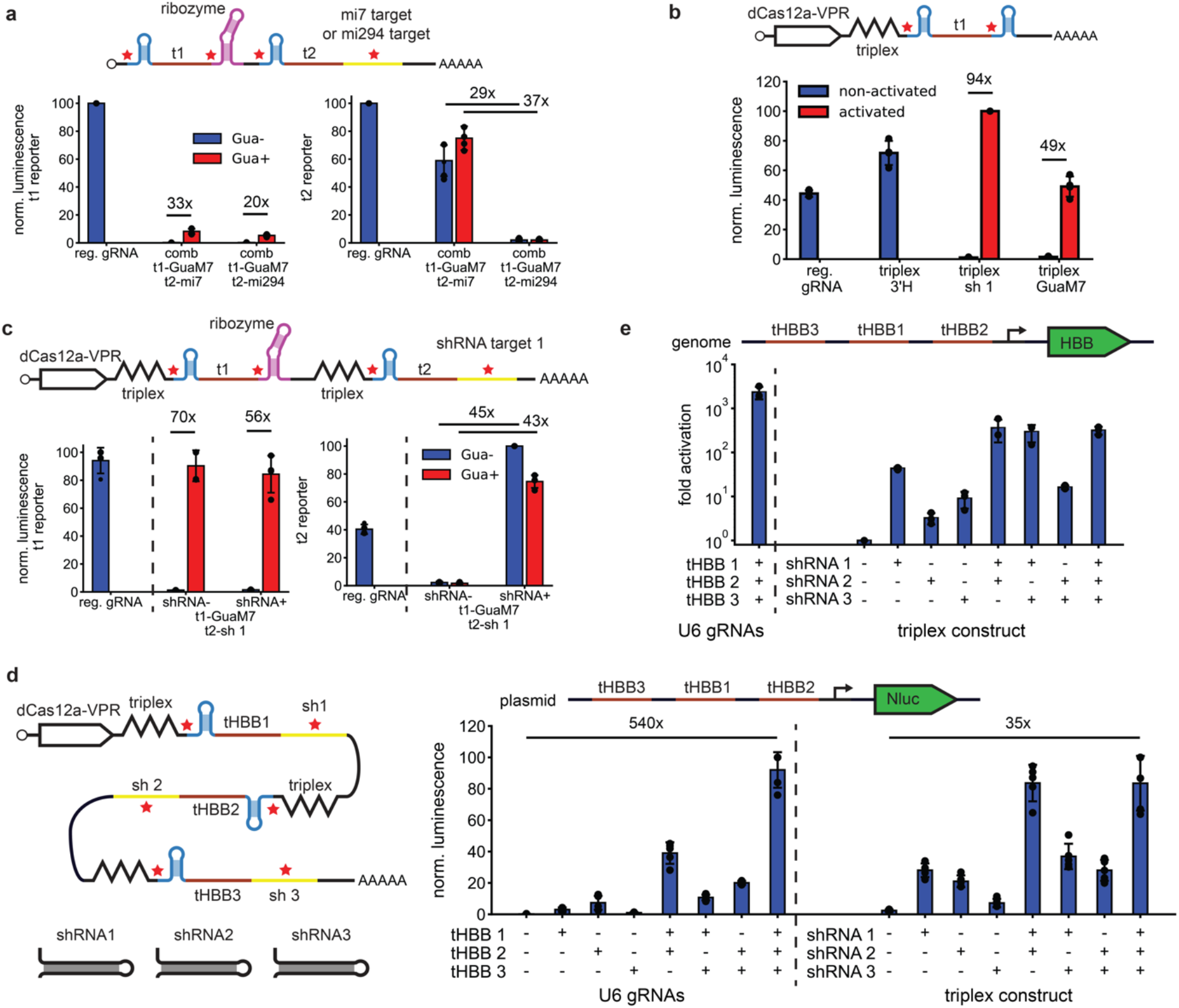
Placing multiple switchable gRNAs on a single transcript. **a** Nanoluciferase assay for two switchable gRNAs placed next to each other on a single CAG promoter transcript. (N=4) **b** Nanoluciferase assay for Cas12a and switchable gRNAs placed on the same transcript and separated using a Malat1 triplex. activated = with shRNA 1 for sh 1 construct, with 100 μM guanine for GuaM7 construct (N=4) **c** Nanoluciferase assay for Cas12a and two switchable gRNAs placed on the same transcript, all of them separated by a Malat1 triplex. (N=4) **d** Nanoluciferase assay for Cas12a and three switchable gRNAs placed on the same transcript, all of them separated by a Malat1 triplex. All gRNAs target the same minimal promoter. (N=5) **e** RT-qPCR measurement of the HBB mRNA level for three switchable gRNAs placed on the same transcript. The fold activation is measured relative to the triplex construct without shRNAs. (N=3)

A similar problem occurs when trying to integrate the Cas12a coding sequence and its gRNAs on a single transcript: after excision of the gRNAs, the mRNA is degraded due to the lack of a poly(A) tail. Campa et al. recently resolved this issue in the context of conventional gRNAs by inserting a protecting RNA triplex structure from the murine Malat1 noncoding RNA between the Cas12a coding sequence and a gRNA array^9^. We tested whether our switching principle was compatible with this approach by separating conditional Cas12a gRNAs from the enAsdCas12a coding sequence in the same manner (Figure 4b). Exhibiting a 94-fold and 49-fold activation by shRNA/RISC or guanine/GuaM7HDV, respectively, the constructs perform even better than our previous gRNA-only constructs.

We reasoned that protection from degradation by insulating RNA triplex structures might also improve the performance of multiple switchable gRNAs arrayed on a single transcript. We therefore placed an additional Malat1 triplex between two switchable gRNAs with a shRNA target site and the GuaM7HDV ribozyme. As desired, the gRNAs on the resulting construct can be activated independently by shRNA and guanine (Figure 4c). A slight reduction in activity for the shRNA-activated gRNA in the presence of guanine is in line with our observations for the wtHDV ribozyme (Figure 2a). The repeated use of the Malat1 triplex therefore appears to be a viable approach to integrate the Cas12a coding sequence along with a switchable gRNA array on a single transcript. We also tested the constructs used in Figure 5b & c in NIH/3T3 mouse fibroblasts (Supplementary Figure 12a & b) to assess whether our approach would work in other cells than HEK293. Both the individual gRNAs and the integrated constructs were found to be functional in the 3T3 cells.

Thus far, our processed gRNAs targeted seven repeats of the same sequence in the nanoluciferase promoter on our reporter plasmid. When targeting endogenous genes, each target sequence will typically only occur once, but transcriptional activation via enAsdCas12a-VPR using only a single binding site tends to be weak. We therefore copied a short region that naturally occurs in the *HBB* (hemoglobin subunit beta) gene in front of our nanoluciferase reporter and picked three Cas12a target sites in this region (Figure 4d).

We then placed the corresponding three gRNAs with three orthogonal shRNA target sites onto a single transcript, representing a type of fuzzy AND gate for the three shRNAs (Figure 4d and Figure 1e). In Figure 4d, we compare the activation achieved by conditional switching with different combinations of the three shRNAs to the activation achieved with conventional U6-transcribed gRNAs. The maximum activation level achieved in the presence of all three gRNAs is the same in both cases, but a larger leak activation results in a lower dynamic range for our conditional gRNA array compared to the U6 gRNAs (35-fold instead of 540-fold). Notably, activation for shRNA1 is much larger than for the corresponding U6 gRNA, and shRNA1 and 2 activate as strongly as all three shRNAs combined. We also compared activation by shRNAs 1 and 2 as compared to all three shRNAs over a range of gRNA dosages, showing that activation by all three gRNAs is significantly better at least in two out of four cases (Supplementary Figure 13).

We finally tested the same conditional gRNA construct for activation of the genomic HBB gene itself (Figure 4e), which we read out via RT-qPCR. The activation is similar as for the Nluc construct, although activation by shRNA 3 is much stronger. Also the activation by a combination of shRNA 1 and 2 or 1 and 3 is much stronger than for any individual shRNA (~500 fold as opposed to ~50 fold for shRNA 1), indicating that all shRNAs activate their target gRNAs as designed. However, activation by all three shRNAs is not stronger than for either of the two combinations shRNA 1/2 or shRNA 1/3, similar as observed with shRNA 1 and 2 for the Nluc reporter.

## Discussion & Conclusion

We have here developed an approach for switchable gene activation via dCas12a-VPR in mammalian cells that is based on the conditional cleavage of Pol II-transcripts containing Cas12a gRNAs. Processing of the transcripts removes any 5’ and 3’ sequence context and thus results in fully active gRNAs, which can be utilized by dCas12a-VPR. Conditional cleavage has been achieved by inserting miRNA/RISC target sites, a co-factor dependent ribozyme, and an auxiliary gRNA handle structure that can be switched with a strand displacement mechanism, resulting in cleavage by Cas12a’s intrinsic RNase activity. We surmise that also other cleavage mechanisms would work in this context in a similar way. Notably, our approach can be extended to integrate multiple conditional gRNAs and even the coding sequence of Cas12a on the same transcript, for which the different sequence domains have to be protected from degradation using RNA triplexes.

While RISC and ribozyme-induced cleavage have been previously employed to exert posttranscriptional control of gene expression, toehold-mediated strand displacement of the pseudoknot structure characteristic of Cas12a gRNAs is a novel switching principle that allowed us to utilize arbitrary input trigger RNA sequences without any sequence constraints. At this point, we still found considerable variability in the performance of our mammalian strand-displacement gRNAs, however, and no clear design-function relationship yet. We expect that improved sequence design that takes into account folding of the complete SD gRNA construct and also the boundary conditions set by the cellular environment such as the presence of the transcriptome and RNA-binding proteins will lead to more predictable behaviors.

Compared to earlier designs of switchable guide RNAs that worked in vitro and in bacteria, our approach geared towards the control of eukaryotic gene expression has two significant advantages: First, the sequence domain responsible for activation of the gRNA is spatially separated and independent from the gRNA sequence itself, which enables a modular combination of multiple control domains in one gRNA transcript. The second advantage of the approach is the compactness of the genetic constructs required for implementing conditional gRNA circuits. All components for the evaluation of potentially quite complex logic expressions can be placed on a single transcript, which can be generated from a single promoter and from a single plasmid. This also considerably simplifies the delivery of such circuits to mammalian cells.

The general approach outlined here is likely to work for Cas9 gRNAs in much the same way. While strand displacement-activated gRNAs are specific to Cas12a, since they are based on handle pseudoknot formation and Cas12a’s RNase activity, aptazyme-based activation should be straightforward to adapt. MicroRNA-activated Cas9 gRNAs were already implemented in earlier work^14^. As shown in the present work, also in this case placing the coding sequence for Cas9/dCas9 on the same transcript as its microRNA-switchable gRNAs could simplify the delivery of a complete conditional gRNA circuit considerably.

In conclusion, we have demonstrated a general approach to design switchable Cas12a gRNAs responding to a wide variety of stimuli via conditional RNA cleavage. The switching principle is compatible with a compact integration of all necessary components on a single transcript, which facilitates the realization of multiplexed gene regulatory circuits for mammalian cells that can be delivered on a single plasmid.

## Acknowledgements

We thank Enikö Baligács for assistance with a variety of tasks in the lab, Dong-Jiunn Jeffery Truong and Gil Westmeyer for providing the initial nanoluciferase plasmid. We thank Gil Westmeyer and Richard Koll for discussions which contributed to the development of the pseudoknot-based strand displacement switching principle. We thank Christoph Gruber for providing HEK293 cells. We thank Jessica Kretzmann for help with the flow cytometry measurements.

This work is funded by the Deutsche Forschungsgemeinschaft (DFG project no. SI761/5-1).

We gratefully acknowledge funding by the European Research Council (ERC grant agreement no. 694410 – project AEDNA) and the Max Planck School Matter to Life.

## Author Contributions

L.O. conceived the work, performed the experiments, and analyzed the data. L.O. and F.C.S. discussed and interpreted the results and co-wrote the paper.

## Competing interests

The authors declare no competing interests.

## Methods

### Cell culture

HEK293 and NIH-3T3 cells were cultured in DMEM (Merck) supplemented with 10% FBS (Merck), 292 μg/ml L-glutamine, 100 U/ml penicillin and 100 μg/ml streptomycin (ThermoFisher) at 37°C in a 5% CO_2_ humidified incubator. Cells were counted using a Countess II automated cell counter (ThermoFisher).

### Cloning

Plasmid sequences can be found in Table 2. DNA oligonucleotide and gBlock (both from IDT) sequences used for cloning can be found in Table 1.

The reporter plasmid containing nanoluciferase under the minimal promoter containing multiple t2 targets was a gift from Dong-Jiunn Jeffery Truong and Gil Westmeyer (HMGU and TU Munich). A reporter plasmid containing multiple t1 target sites was constructed by restriction digestion of the original reporter plasmid with SfiI digestion and overlapping DNA oligos. The overlapping DNA oligos containing t1 target sites were designed to form a ladder with different numbers of repeats upon ligation. The correct band was excised from a polyacrylamide gel and used in the ligation reaction. The fluorescent reporter plasmid was created from the original nanoluciferase plasmid using Gibson assembly. The reporter plasmid containing the HBB gene target sites was constructed using SfiI digestion and a gBlock.

The enAsdCas12a-VPR activator plasmid was obtained from Addgene (https://www.addgene.org/107943/) ^3^. U6-promoter plasmids for expression of gRNAs were based on the BPK3079 plasmid, which already contains Esp3I sites (https://www.addgene.org/78741/)^32^. For the expression of triggers from the U6 promoter, the Cas12a handle sequence of the BPK3079 plasmid was removed using overhang PCR. A CAG promoter plasmid with two Esp3I sites was created via restriction-digestion of the dCas12a-VPR plasmid with EcoRI and NotI and overlapping oligos.

Most constructs for gRNAs or strand displacement triggers were assembled using GoldenGate cloning in combination with overlapping oligos. DNA oligos were purchased from IDT and annealed at a concentration of 2 μM in H_2_O with 2 mM MgCl_2_ by cooling them from 90°C to 20°C over a period of 20 min. For the GoldenGate reactions, 3 nM of plasmid, 15 nM of each overlapping oligo pair, 1 U/μl Esp3I (NEB), 0.5 U/μl T4 Polynucleotide Kinase (NEB), and 40 U/μl T4 DNA Ligase (NEB) were mixed in T4 DNA Ligase Reaction Buffer (NEB). The mix was cycled between 37°C for 60s and 16°C for 90s fifteen times, kept at 37°C for 10 min, deactivated at 70°C for 10 min and chemically transformed into DH5alpha.

The combination plasmids in Figure 5a were obtained from the original CAG-gRNA plasmids by using overhang PCR to introduce Esp3I restriction sites and GoldenGate cloning as described above. The dCas12a-VPR-triplex-gRNA-t1 plasmid was assembled using Gibson assembly with a gBlock after digestion of the dCas12a-VPR plasmid with NotI. The dCas12a-VPR-triplex-gRNA-t1-sh1/GuaM7HDV plasmids were obtained using restriction-digestion with XhoI and NotI, the sites for which were inserted into amplicons from gRNA-only plasmids using overhang PCR. Triplex plasmids containing multiple gRNAs were created using Gibson assembly and gBlocks.

Plasmids were transformed using chemical transformation. Chemically competent cells (40 μl) were thawed on ice. 5 μl of ligation reaction plasmid were added to the cells, the liquid was mixed, and incubated on ice for 20-30 min. The cells were placed at 42°C for 45s in a water bath and placed back on ice. 950 μl of LB medium were added and the mixture was incubated at 37°C for 20-30 min. The cells were plated on LB agar with suitable antibiotics and left in an incubator at 37°C overnight. Colonies were picked, diluted in LB medium, incubated overnight and plasmids were purified using ZymoPURE Plasmid Miniprep Kits according to the manufacturer’s instructions. The plasmids were sequenced using the primers listed in Table 1.

### Plasmid transfection

We used Xfect transfection reagent (TakaraBio) according to the manufacturer’s instructions to transfect plasmids. We seeded approximately 160000, 80000, and 45000 cells per well in 12-well (Nunc Cell-Culture Treated, ThermoFisher), 24-well (VWR Tissue Culture Plates, Surface Treated), and 48-well (CELLSTAR, Greiner Bio-One) plates in a total volume of 600 μl, 300 μl, and 180 μl 24 hours before transfection, respectively. A total of 2000, 1000, or 600 ng of plasmids (for 12-well, 24-well, and 48-well plates) were added together for transformation of a single well and filled up to 50 μl, 25 μl, and 15 μl with Xfect reaction buffer. When the total DNA content was less than these values, the rest was added using a filler plasmid. 0.6 μl, 0.3 μl, and 0.18 μl of Xfect transfection reagent were added and the tubes were vortexed for 10 s at maximum speed. The liquid was collected at the bottom by tapping the tubes on a table surface and the mixture was incubated for 10 min at RT. The transfection mix was added dropwise to the wells and the cultures were incubated for 4 h. The old medium was removed and 600 μl, 300 μl, and 180 μl of fresh medium were added. Inducers (i.e., guanine) were added at this point if necessary. The transfected cells were cultured for two days before luciferase measurements, fixation for microscopy, or two to three days for RNA purification.

### Luciferase measurements

Transfections for luciferase measurements were carried out in 48-well plates. For experiments with separate dCas12a-VPR and gRNA plasmids and without RNA triggers, we used 300 ng of nanoluciferase reporter plasmid, 150 ng of dCas12a-VPR plasmid, 66 ng of each U6-gRNA plasmid, 150 ng of each CAG-gRNA plasmid, and 5 ng of shRNA plasmid (unless specified otherwise). For experiments with separate dCas12a-VPR and gRNA plasmids and with RNA triggers, we used 240 ng of nanoluciferase reporter plasmid, 120 ng of dCas12a-VPR plasmid, 60 ng of each CAG-gRNA plasmid and 180 ng of each trigger plasmid (unless specified otherwise). For the experiments with the Malat1 triplex-based design, we used 300 ng of nanoluciferase reporter plasmid, 300 ng of dCas12a-gRNA plasmid, and 6 ng of each shRNA plasmid.

Two days after transfection, we sampled the supernatant. For HEK293, we diluted 20 μl of supernatant in 80 μl of DMEM. For NIH/3T3 cells, we used undiluted supernatant. A master mix containing 0.2 μl of Nano-Glo Luciferase Assay Substrate (Promega), 9.8 μl Nano-Glo Luciferase Assay Buffer, and 5 μl PBS per sample was prepared. The supernatant was added to the side of PCR tubes and the master mix was added. Sample and master mix were mixed by centrifuging the strip in a table centrifuge. The mix was transferred to a Nunc white 384-well plate (ThermoFisher) using a multichannel pipette. The luminescence was measured in a CLARIOstar microplate reader (BMG Labtech). To account for variations in total luciferase intensity due to differences in the transfection reagent, cell number, or other factors, we normalized each experiment to the highest total luciferase value occurring in that experiment.

### Flow cytometry

Transfections were carried out in 24-well plates. We used 500 ng of mScarlet reporter plasmid, 250 ng of dCas12a-VPR plasmid, 110 ng of each U6-gRNA plasmid and 250 ng of each CAG-gRNA plasmid. After two days, cells were detached with 100 ul of 0.05% Trypsin-EDTA for 5 min at 37°C. 300 ul of DMEM and 400 ul of 4% formaldehyde solution was added. The cells were fixed for 10 min at RT and spun down at 350 g for 5 min. The supernatant was removed, the cells were resuspended in 1 ml of PBS, and kept at 4°C until the measurement.

Samples were acquired and analyzed using an AttuneTM NxT Flow Cytometer and software (ThermoFisher) respectively. 20,000 single cell events, gated on side scatter area vs height, were recorded for analysis. mScarlet was excited by a 561 nm laser, and emission was measured with a 620/15 nm bandpass filter. We used FlowJo V10.1 to analyze the data. Around 70-80% of cells were contained within the preliminary FSC-SSC gating. The bar graphs displayed in the figures show the average fluorescence of the gated cell population normalized to the highest average fluorescence occurring in the experiment.

### RT-qCPR

Transfections were carried out in 12-well plates. For Figure S11, we used 440 ng of the U6-gRNA plasmids, 1000 ng of the CAG-gRNA plasmids, 1000 ng of dCas12a-VPR with a suitable amount of fill plasmid to bring the total amount to 2000 ng. RNA was harvested two days after transfection. For Figure 5e, we used 333 ng of each U6 gRNA plasmid and 1000 ng of dCas12a-VPR plasmid or 1910 ng of the dCas12a-gRNA-triplex construct and 30 ng of each shRNA. For Figure S11, RNA was harvested two days after transfection. For Figure 4e, RNA was harvested three days after transfection. Total RNA was purified using Quick-RNA Miniprep Kits (Zymo Research). RNA concentration and quality was assessed using a NanoPhotometer (Implen). Reverse transcription was performed using M-MulV Reverse Transcriptase (NEB) with 250 ng of total RNA using gene specific primers for Figure S11 and using an oligo-T primer for Figure 4e. After reverse transcription, the cDNA was diluted 1:6 in nuclease-free H_2_O. qPCR was carried out using the primers in Supplementary Table 3 and Luna Universal qPCR Master Mix (NEB) in a BioRad iCycler in technical triplicates for each biological replicate. The program for all sets of primers except for the quantification of gRNA was 95°C for 60s, 35 or 45 cycles of 95°C for 15s and 60°C for 30s. For gRNA quantification, the program was 95°C for 60s, 35 cycles of 95°C for 15s, 53°C for 15s, and 60°C for 30s. Efficiency data were obtained from calibration curves. Expression data were normalized to the level of ACTB expression. RNA was quantified using the efficiency-corrected delta-delta Cq method.

### Fluorescence in-situ hybridization (FISH)

Transfections were carried out in 24-well plates. We used 220 ng of the U6-gRNA plasmids, 500 ng of the CAG-gRNA plasmids, 500 ng of dCas12a-VPR with a suitable amount of fill plasmid to bring the total amount to 1000 ng. One day after transfection, the cells were detached from their well using Trypsin-EDTA. The suspension was diluted to a total of 1 ml of liquid per well using DMEM, of which 400 μl were transferred to a new well which was treated with poly-D lysine to improve adherence during washing steps. The next day, the medium was removed and cells were fixed for 10 min in 4% formaldehyde solution (Merck). The fixation solution was removed and the cells were washed twice with nuclease-free H_2_O (nfH_2_O). The cells were permeabilized by incubation in 70% ethanol overnight at 4°C. The ethanol was removed and the cells were incubated in 300 μl of wash buffer (2x SSC, 10% formamide) for 5 min at room temperature. A total of 150 μl of hybridization solution was added to each well (200 nM Cy5-labeled FISH probe, 2x SSC, 10% formamide, 10 wt% dextran sulfate). The mixture was incubated at 37°C for 4 h. Afterwards, cells were first washed with 300 μl wash buffer, then incubated with wash buffer at 37°C for 15 min. Nuclei were stained by incubating the cells in wash buffer supplemented with 1 μg/ml Hoechst 33342 (ThermoFisher) for 10 min at room temperature. Cells were imaged in 2X SSC buffer using an EVOS M7000 cell culture microscope (ThermoFisher) using 20x magnification. Hoechst 33342 was measured using an excitation and emission of 357/44 nm and 447/60 nm. Cy5 was measured using an excitation and emission of 628/40 nm and 685/40 nm. Data were evaluated using ImageJ.

### In vitro transcription template generation

DNA template sequences for in vitro transcribed RNA were constructed by adding a T7 promoter in front of the designed RNA sequences. When the secondary structure allowed it, the resulting template sequences were split into two parts with sufficient overlap. The two parts were ordered as singlestranded DNA oligos from Integrated DNA Technologies (IDT). Alternatively, the entire sequence was ordered as an Ultramer from IDT and a T7 promoter primer was used for the following protocol. The double-stranded template was produced by annealing the strands and filling in the single-stranded regions with Phusion HF PCR Master Mix (NEB) by cycling between the melting temperature of the strands and 72°C twenty times. mCerulean and mCherry templates were PCR amplified from plasmids. The resulting DNA templates were column purified using a Monarch PCR&DNA Cleanup Kit (NEB) and quantified by their absorption on a NanoPhotometer (Implen). The length of the templates was verified in a native 12% polyacrylamide (29:1 acrylamide/bis-acrylamide) gel run at 120 V for 45 min by comparing with a dsDNA ladder (Low Molecular Weight DNA Ladder, NEB).

### In vitro transcription

The transcription mixture with a total volume of100 μl contained approximately 80 nM of DNA template, 1x T7 RNA Polymerase buffer (80 mM Tris-HCl, 6 mM MgCl_2_, 1 mM DTT, 2 mM spermidine, pH 7.9), 4 mM of each rNTP, 12 mM MgCl_2_ and 1 U/ μl T7 Polymerase (all NEB). The transcription mix was incubated at 37°C for 4 h. Afterwards, 3 units of DNaseI (NEB) were added and the mixture was incubated at 37°C for 1 h.

The RNA was gel purified. It was run on an 8 M urea denaturing polyacrylamide (29:1 acrylamide/bis-acrylamide) gel in 1x TBE buffer. The percentage of polyacrylamide was chosen based on the length of the RNA to be purified and tended to be between 6% for transcribed mRNAs with a length of roughly 1000 nt and 12% for the shortest RNAs used. The gels were first heated to 65°C using a JULABO water circulator while being run empty at 30 V for 30 min. The samples were mixed 1:1 with Gel Loading Buffer II (ThermoFisher) and incubated at 90°C for 5 min. The samples were then loaded onto the gel and run at 120 V for 45 min. They were stained for 15 min with SYBR Green II (ThermoFisher), and excised on an illuminating table with blue light. The gel slice was then purified using a ZR small-RNA PAGE Recovery kit (Zymo Research) according to the manufacturer’s instructions.

The length of the purified RNA was then checked in a gel prepared as described in the preceding paragraph. To check the length, we used either a DNA ladder (Low Molecular Weight DNA Ladder or 1 kb Plus DNA Ladder, NEB) or an RNA ladder (RiboRuler Low Range or High Range RNA Ladder, ThermoFisher). The RNA ladders were used to quantify RNA concentration by comparing the total intensities of the bands of purified RNA with the total intensities of the ladder bands of a known concentration using ImageJ.

### In vitro ssDNase assays

We used a short reporter DNA oligo with the sequence FAM-TTATT-BHQ1^33^. A single reaction had a total volume of 15 μl and was run with 1x NEBuffer 3.1, 7 nM of each target, 100 nM reporter oligo, 60 nM of AsCas12a, 17 nM of SD gRNA, and 33 nM of the sensed RNA. The components were assembled at 4°C, transferred into white tubes with clear caps (ThermoFisher) and measured at 37°C in a BioRad iCycler for 1 h.

The data were treated as follows: The baseline was subtracted by shifting the lowest fluorescence value for all samples to zero. The fluorescence was normalized by dividing all fluorescence curves by their highest value if the reaction finished within the timeframe of the experiment or by an average of the other finished values if the reaction did not finish within the timeframe of the experiment. For bar graphs, the fluorescence value 20 min after the start of the experiment was determined and used as the measured value for that bar graph.

### Statistical Analysis

The number of biological replicates is given in figure legends. Error bars show the sample standard deviation. When statistical significance is indicated, it is based on a two-tailed t-Test with alpha=0.05.

## Data availability

All the conclusions drawn in this paper are derived from data shown either in the main text or in the Supplementary Information. All data and sequences are available from the corresponding author upon reasonable request.

## Supplementary Information

**Supplementary Figure 1.**
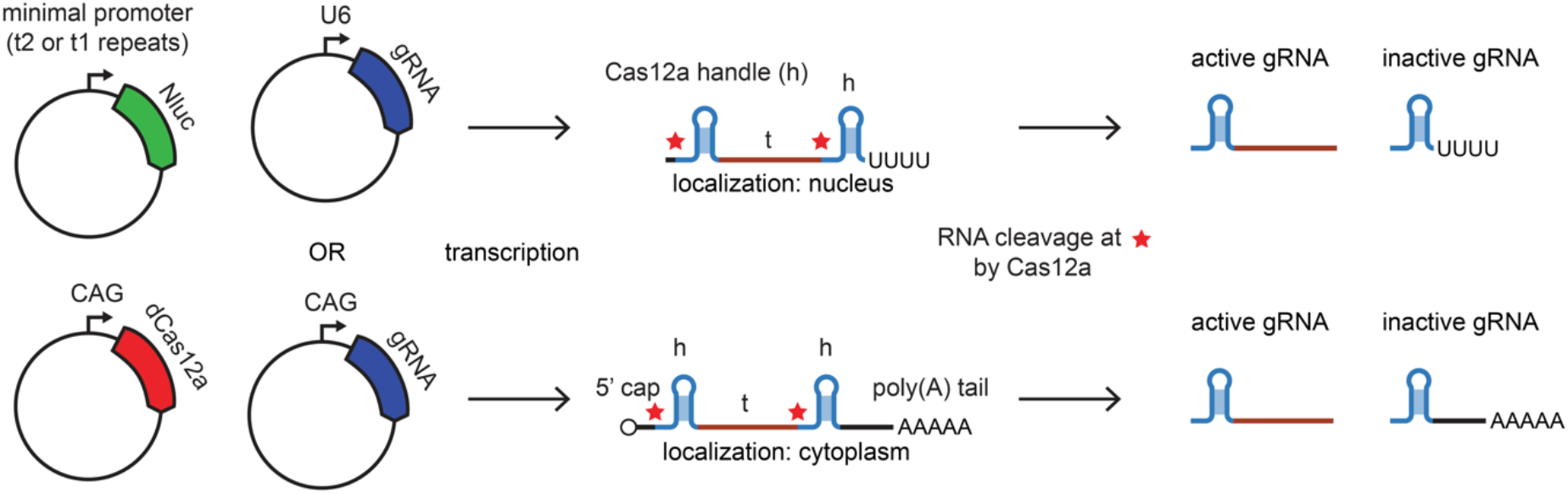
Plasmids used for nanoluciferase measurements and the expected fate of U6- and CAG-transcribed gRNAs.

**Supplementary Figure 2.**
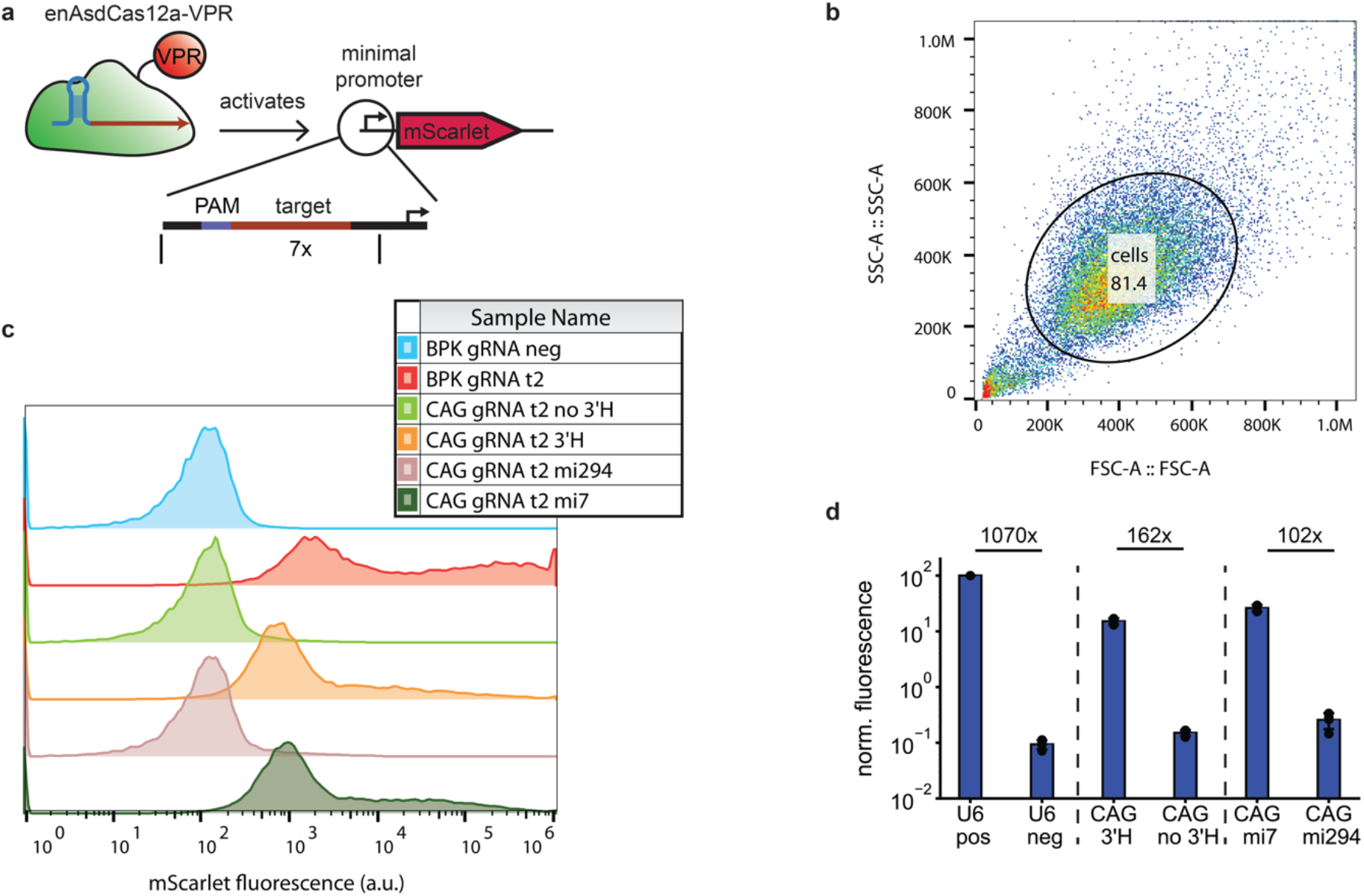
Flow cytometry data for microRNA-activated CAG gRNAs. **a** The mScarlet reporter contains seven repeats of the t2 target sequence. **b** Example of the cell population gating using FSC and SSC. **c** Distribution of fluorescence intensities for different U6 (BPK)- and CAG-transcribed gRNAs. **d** Average fluorescence intensities of populations shown in c. pos = t2 spacer (N=4)

**Supplementary Figure 3.**
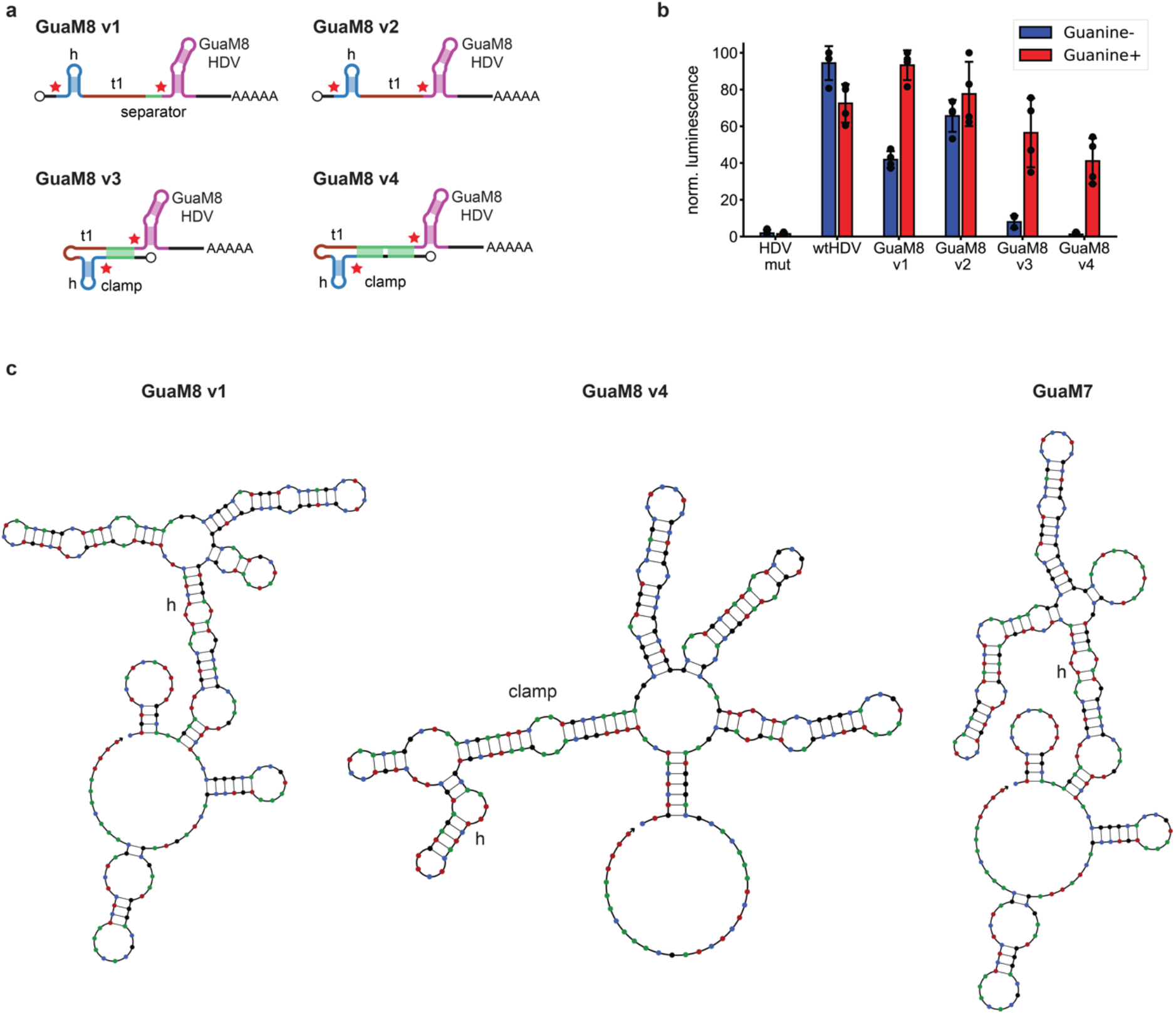
Optimization of GuaM8HDV gRNAs. **a** Design of different GuaM8HDV gRNAs (versions v1 to v4, h = handle, t1 = gRNA spacer). **b** Nanoluciferase assays of the gRNA activity for four different versions of the GuaM8HDV gRNA. (N=4) **c** NUPACK prediction of the MFE structure for different gRNA-ribozyme combinations including 31 nt of the transcript sequence to the 5’ of the Cas12a handle.

**Supplementary Figure 4.**
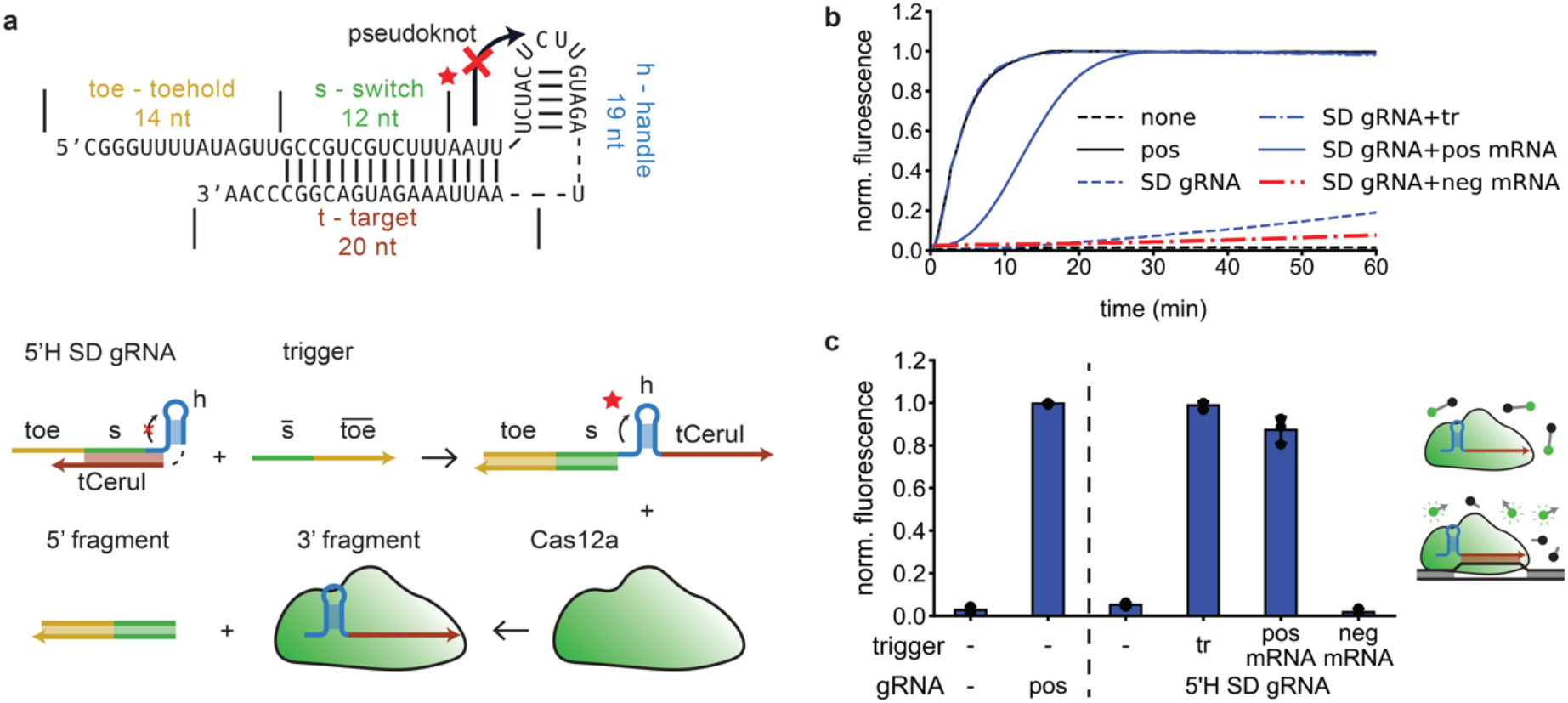
Design of an in vitro strand displacement gRNA based on pseudoknot switching. **a** A 5’ switch domain inhibits pseudoknot formation. Addition of a trigger allows for formation of the pseudoknot, allowing Cas12a to bind and cleave off the 5’ switch. The SD gRNA has a hairpin at the 5’ end and the trigger has hairpins at the 5’ and 3’ to help with transcription^19^. For clarity, the hairpins are omitted in the illustrations. **b** ssDNase assay of the activation of the 5’H SD gRNA by a trigger containing only the sensed sequence (tr), an mRNA containing the sensed sequence (pos mRNA, mCerulean), and a negative mRNA without the sensed sequence (neg mRNA, mCherry). pos = regular gRNA **c** Fluorescence at 20 min for measurements as shown in b. (N=3)

**Supplementary Figure 5.**
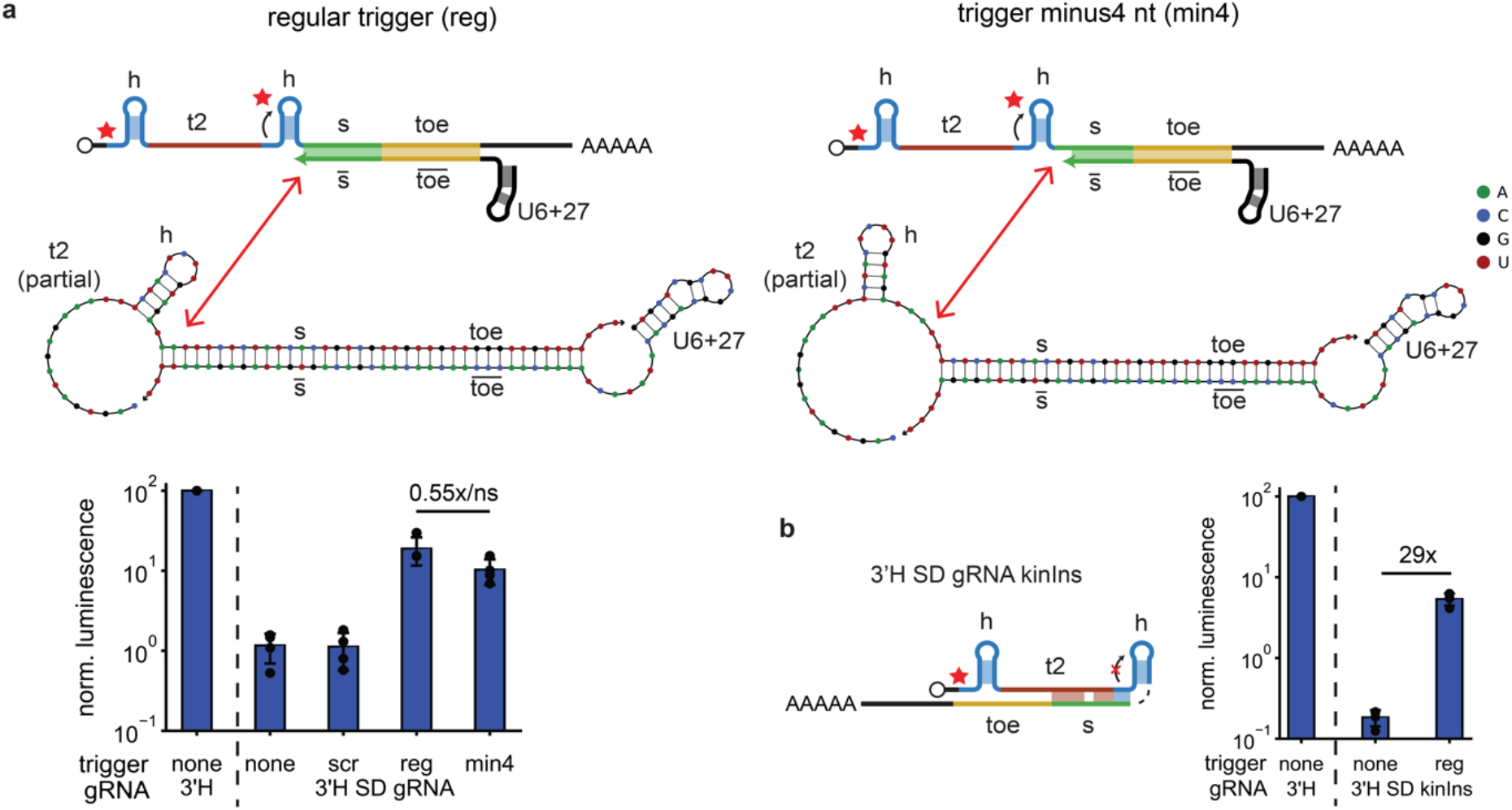
Removal of trailing nucleotides and a kinetic insulator design for 3’H SD gRNAs. **a** Change in trigger activity when the last four nucleotides of the trigger are removed (min4). The red arrows compare schematic representations with NUPACK simulations. reg = trigger with fully complementary sequence, scr = scrambled trigger (N=4) **b** Nanoluciferase assay of a 3’H SD gRNA containing kinetic toeholds with and without trigger. reg = trigger with the fully complementary sequence (N=4)

**Supplementary Figure 6.**
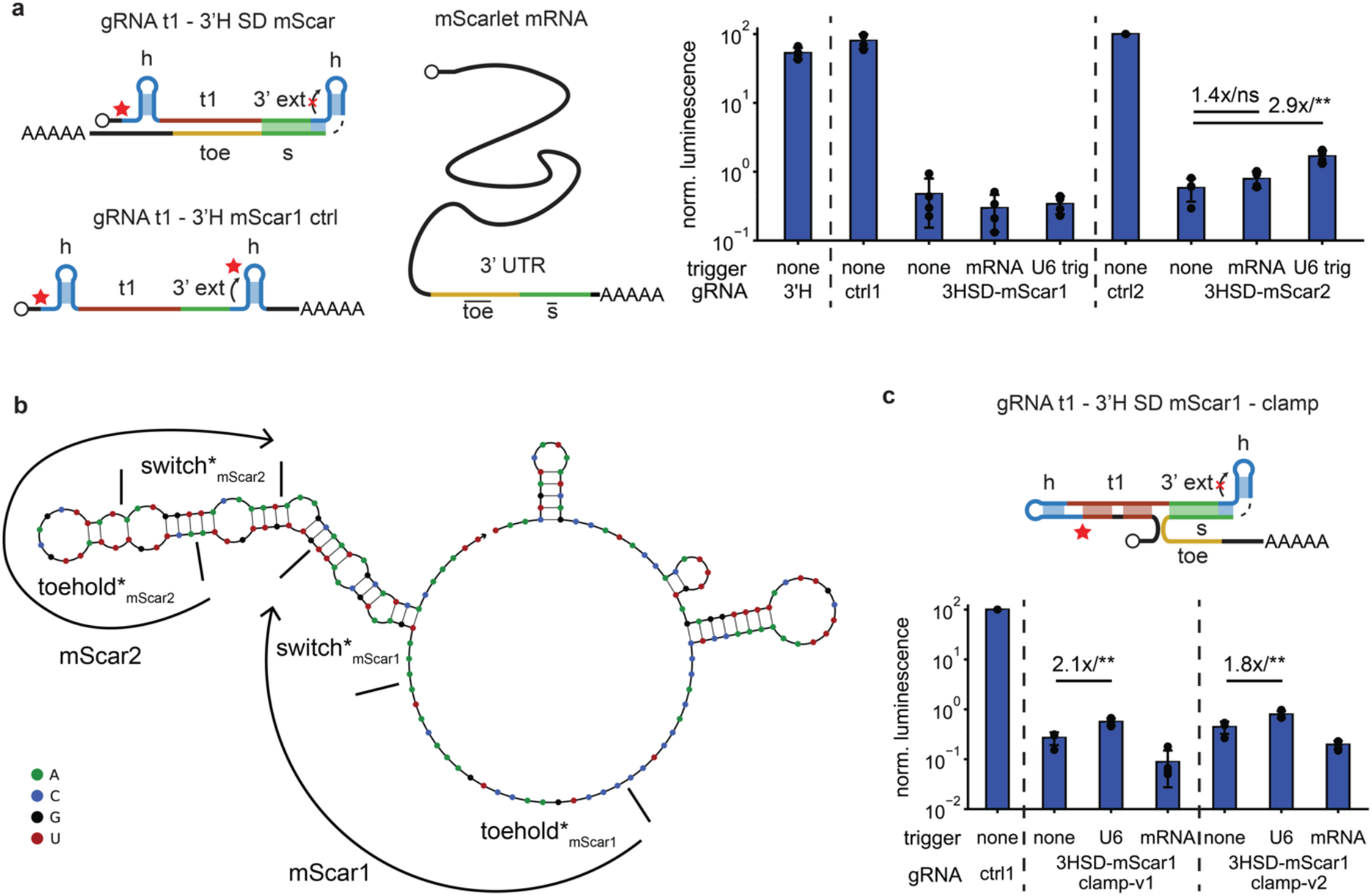
Sensing natural RNA sequences using 3’H SD gRNAs. **a** Nanoluciferase assays of two different 3’H SD gRNAs sensing different sequences on an mScarlet mRNA. 3’H = gRNA with regular 3’ handle, ctrl1/2 = gRNA containing the same 3’ extension before the 3’ handle, mRNA = addition of a plasmid containing mScarlet under a CAG promoter, U6 trig = addition of a plasmid containing only the sensed subsequence under a U6+27 promoter. (N=4) **b** Illustration of sensed subsequences on the mScarlet mRNA 3’ UTR. **c** Clamping the gRNA spacer sequence increases the activation by mScarlet trigger 1. (N=4)

**Supplementary Figure 7.**
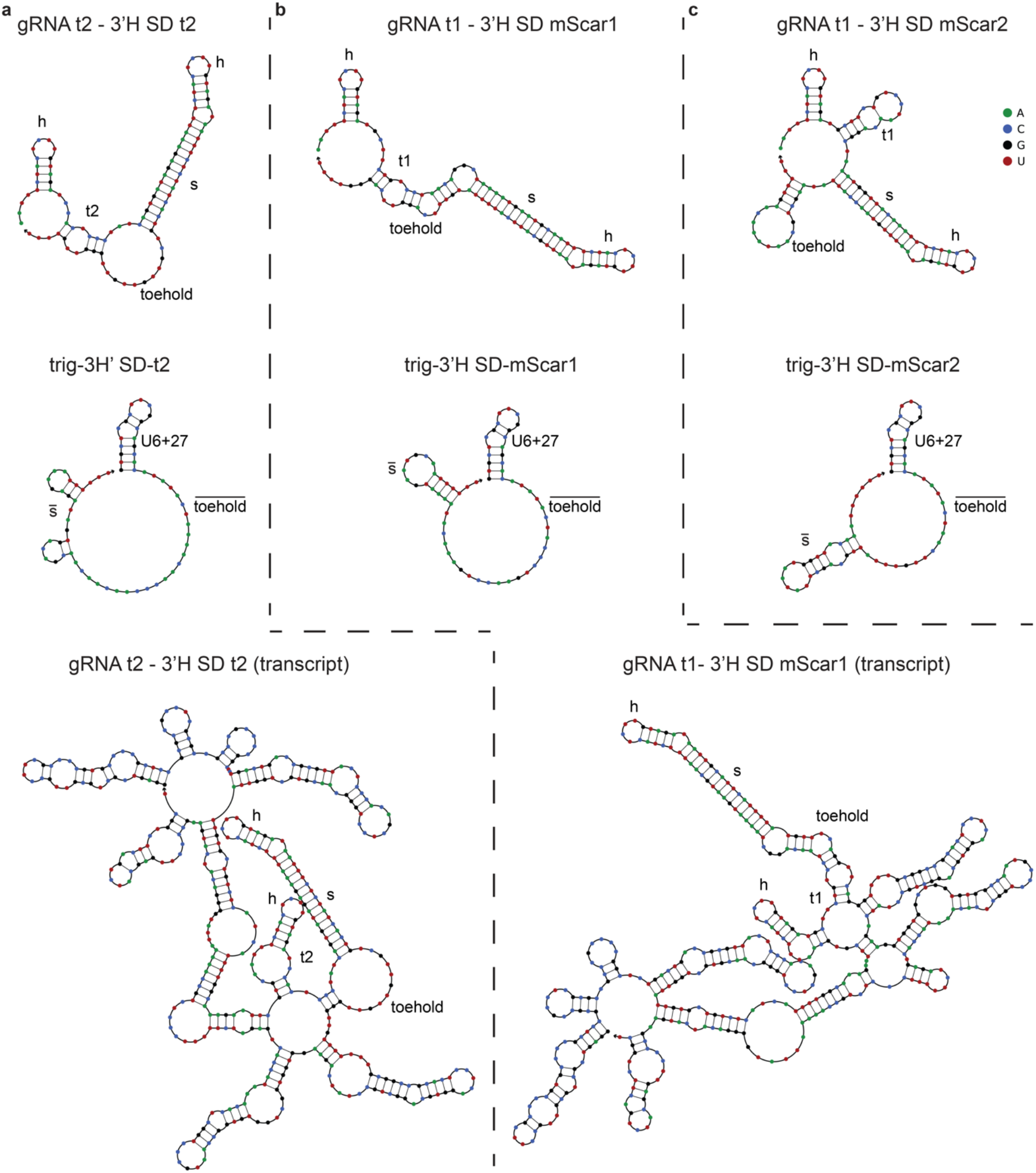
Secondary structure of 3’H SD gRNA designs. NUPACK MFE structure prediction for 3’H SD gRNAs, their associated U6+27 triggers and of the expected transcript sequence for **a** the artificial t2 switch, **b** mScarlet sequence 1, and **c** mScarlet sequence 2. h = handle; t2, t1 = gRNA spacer sequences; s = switch

**Supplementary Figure 8.**
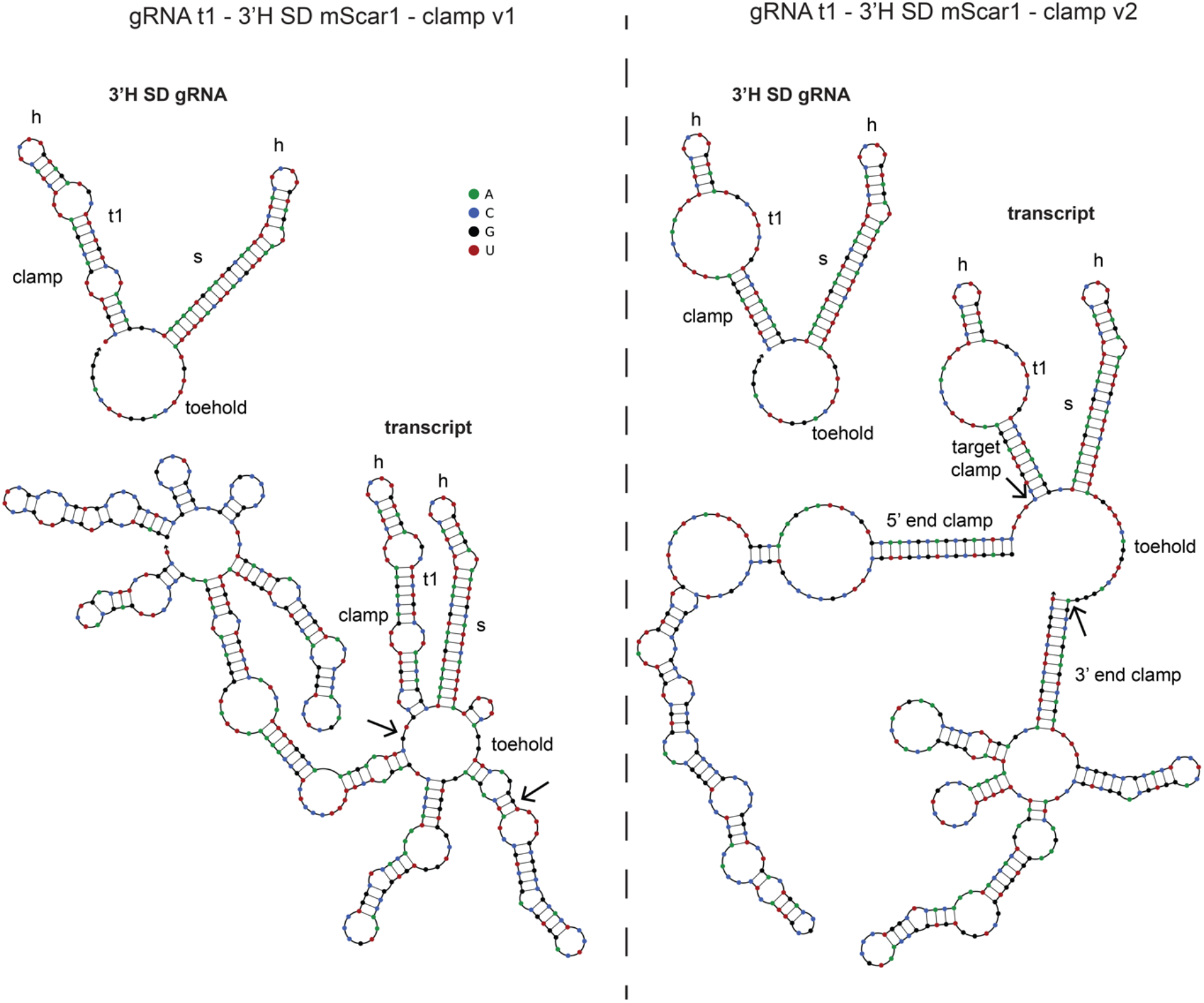
Secondary structure of 3’H SD mScarlet-3’ UTR-sensing designs with clamps. Both the functional gRNA subunit as well as the expected transcript are shown. The arrows indicate the core 3’H SD gRNA sequence. Simulations were performed using NUPACK.

**Supplementary Figure 9.**
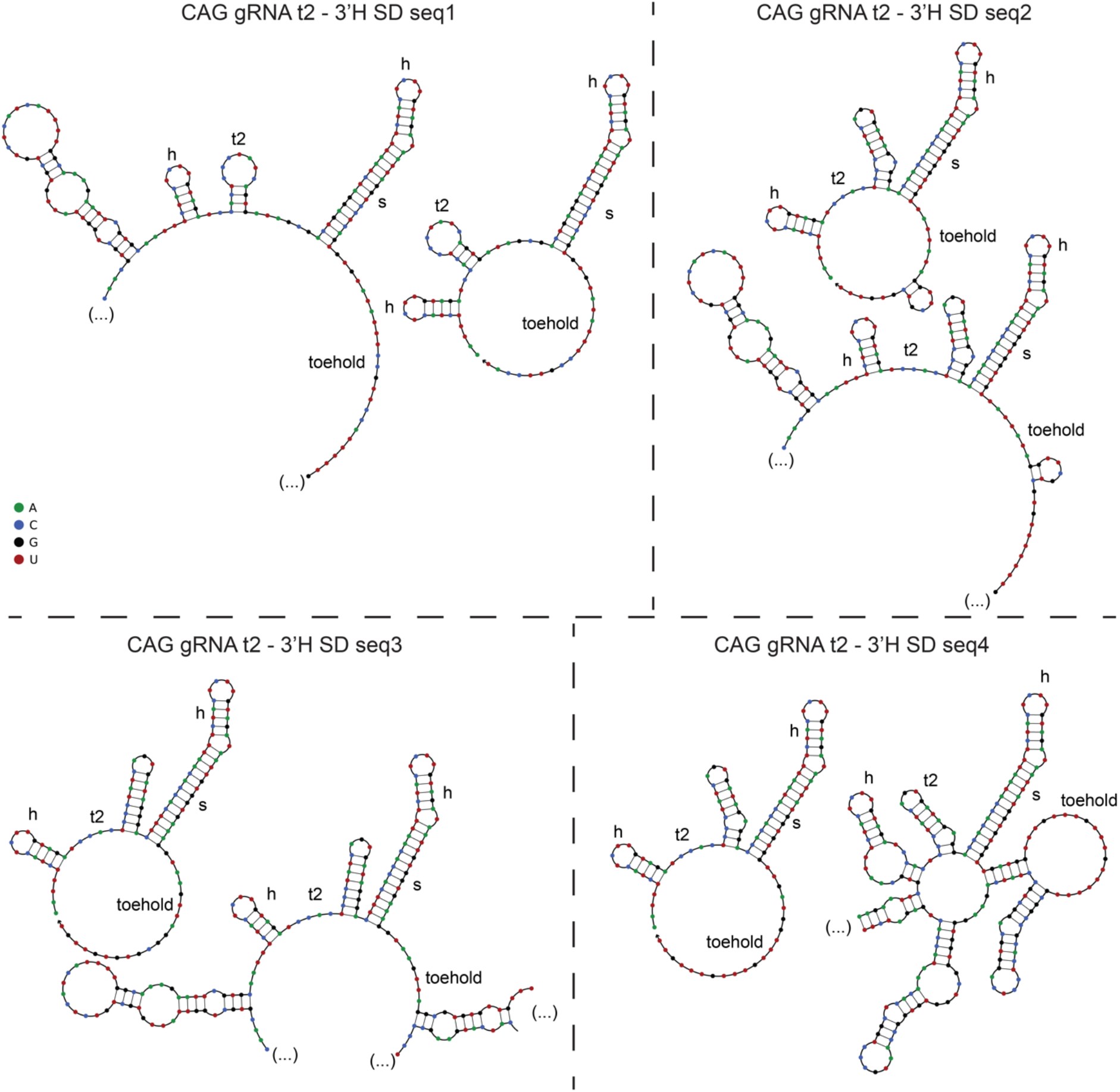
Secondary structure of 3’H SD gRNAs sensing de-novo designed sequences. The functional 3’H SD gRNA and an excerpt of the full expected transcript are shown. Simulations were performed using NUPACK.

**Supplementary Figure 10.**
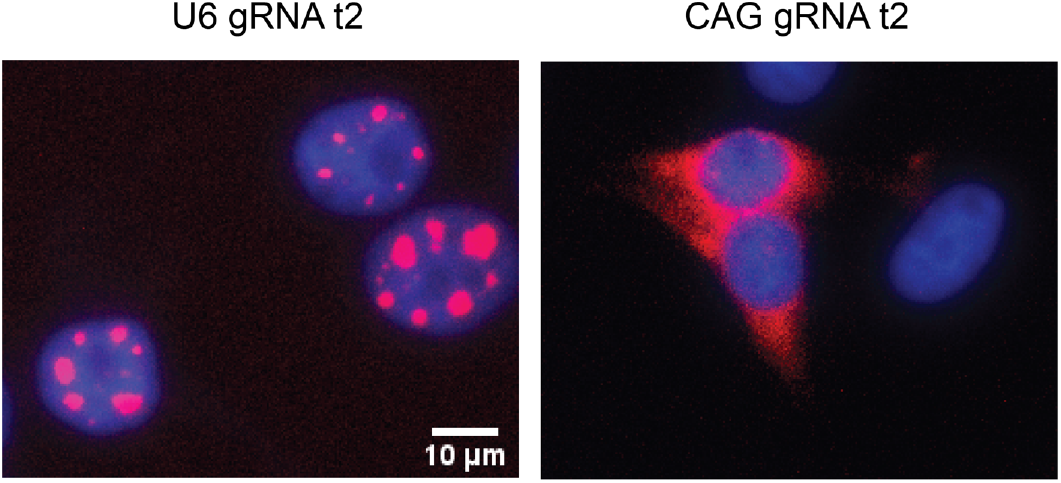
Localization of U6 and CAG-transcribed gRNAs. Fluorescence in-situ hybridization (FISH) experiments targeting the gRNA spacer t2 for U6- and CAG-transcribed gRNAs in the absence of a Cas12a plasmid. The FISH probe (Cy5) is labeled in red and the nucleus (Hoechst 33342) is labeled in blue. U6-transcribed gRNAs are primarily localized in discrete nuclear loci, while CAG-transcribed gRNAs are primarily localized in the cytoplasm.

**Supplementary Figure 11.**
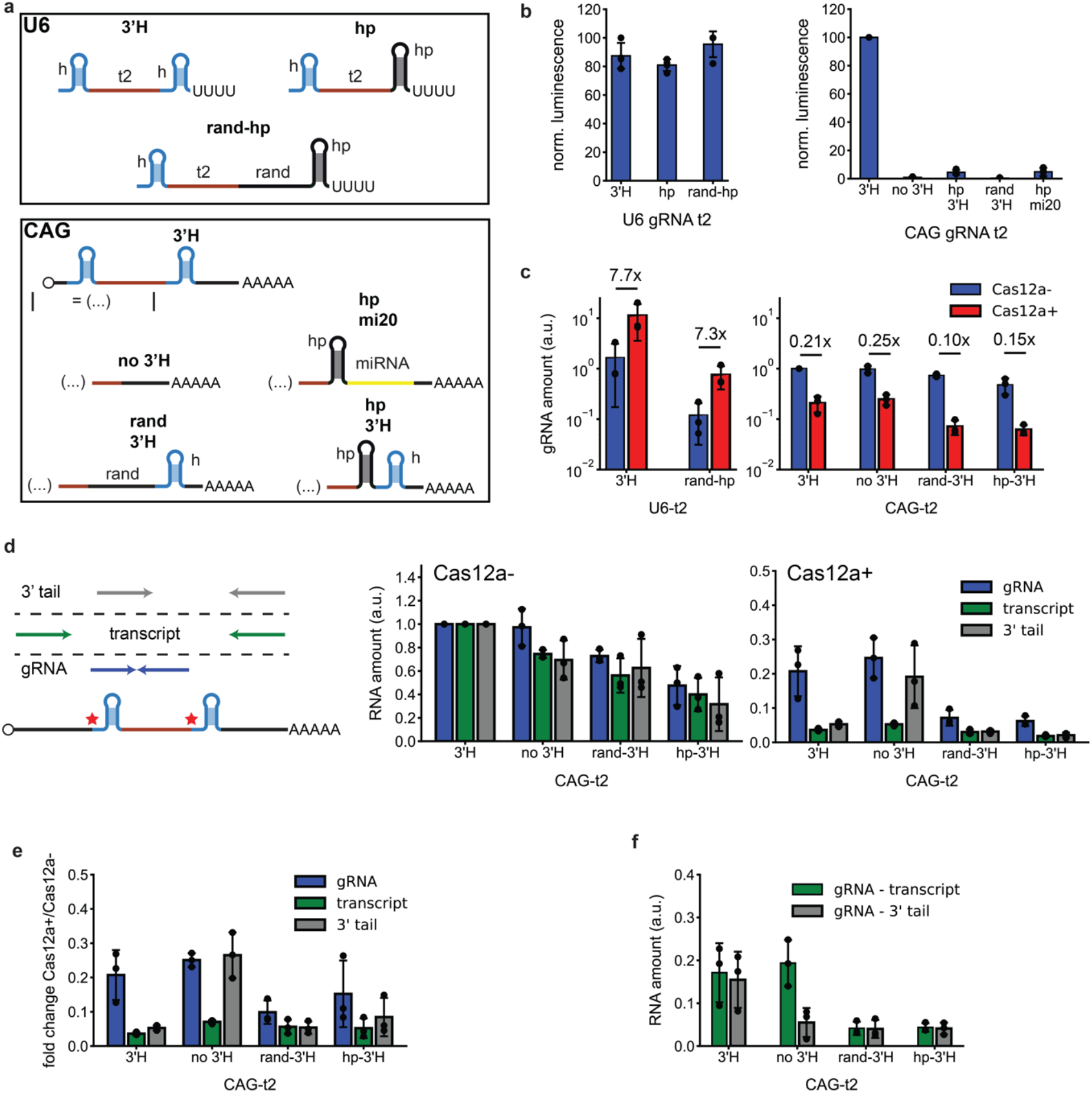
The influence of the 3’ sequence on the switching principle. **a** Different 3’ extensions of U6-transcribed and CAG-transcribed gRNAs. The (…) denotes the 5’ end of the CAG-transcribed gRNAs, containing a 5’ cap, a Cas12a handle and the spacer. h = Cas12a handle, hp = hairpin, rand = randomly generated sequence **b** Luciferase assay of the activity of different extended U6- and CAG-transcribed gRNAs. (N=4) **c** RT-qPCR data of the amount of gRNA with and without the presence of Cas12a for different extended U6- and CAG-transcribed gRNAs. (N=3) **d** RT-qPCR data for the amount of non-processed transcript, processed transcripts retaining a 3’ tail and the gRNA amount without and with addition of the Cas12a plasmid. (N=3) **e** Ratio between the amounts of RNA with and without the Cas12a plasmid for the data shown in d. (N=3) **f** Difference between the gRNA amount and the non-processed transcript or 5’ processed transcript amount shown in d. (N=3)

**Supplementary Figure 12.**
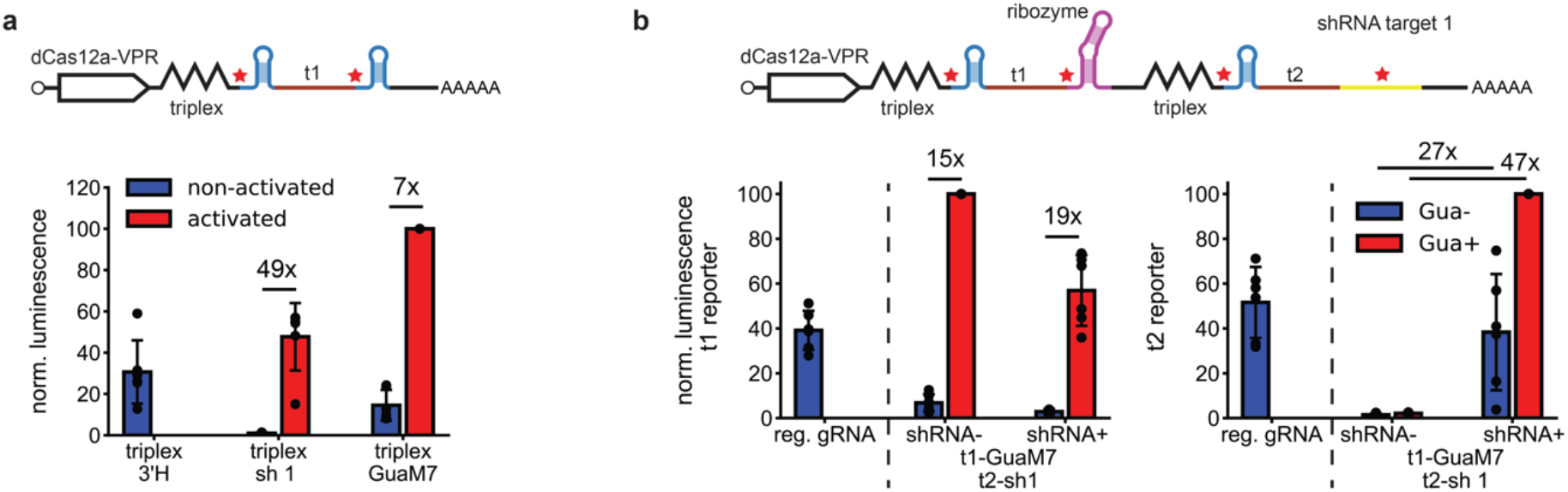
Functionality of the triplex-based dCas12a-gRNA design in NIH/3T3 cells. **a** Nanoluciferase assay for Cas12a and switchable gRNAs placed on the same transcript and separated using a Malat1 triplex. activated = with shRNA 1 for sh 1 construct, with 100 μM guanine for GuaM7 construct (N=6) **b** Nanoluciferase assay for Cas12a and two switchable gRNAs placed on the same transcript, all of them separated by a Malat1 triplex. reg. gRNA = single gRNA with 3’ handle under CAG promoter (N=6)

**Supplementary Figure 13.**
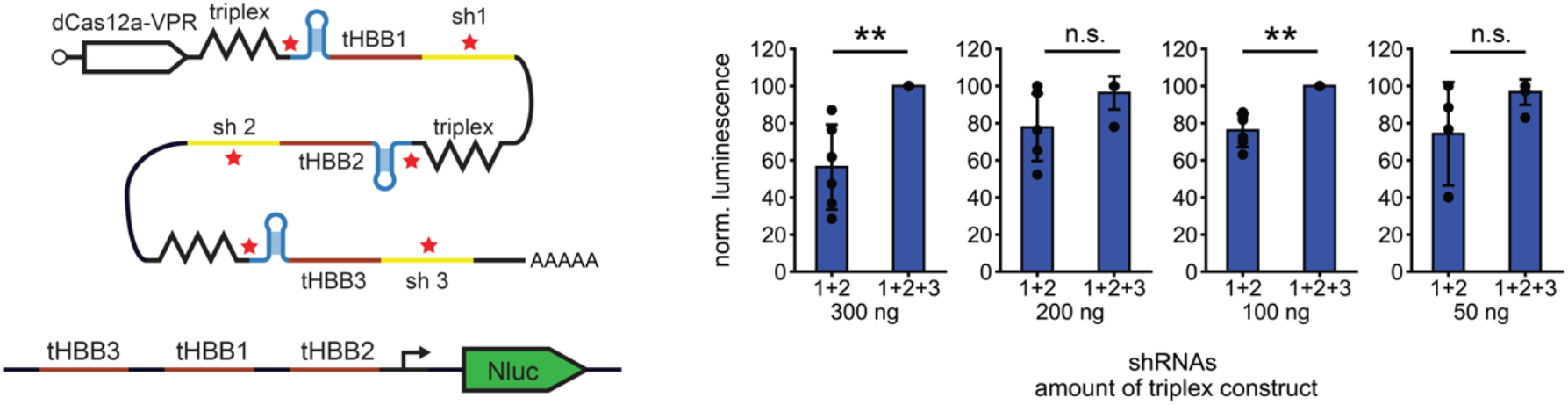
Activation of a multiplexed dCas12a-gRNA construct by two and three shRNAs. **a** Nanoluciferase assay for Cas12a and switchable gRNAs placed on the same transcript and separated using a Malat1 triplex. The total amount of triplex construct was varied. For constructs with less total Cas12a construct, more nanoluciferase plasmid was used. (N=6)

## Supplementary Note 1 – Ribozyme optimization for mRNA inactivation and gRNA activation

As mentioned in the discussion of the results of Figure 2, the disparity between the performance of the GuaM8HDV and GuaM7HDV for mRNAs and our Pol II-transcribed gRNAs could be explained by different optimization objectives for mRNA *inactivation* and gRNA *activation*.

For mRNA inactivation, levels of the measured fluorescent proteins will be approximately linear in the amount of mRNA at the steady state. If the mRNA is produced at a rate P, has a natural decay rate of k_nat_ (with an associated half-life T_nat_=ln2/ k_nat_) and a decay rate due to the ribozyme of k_+/-_ (+/-: in presence/absence of ligand) with an associated half-life T_+/-_, the change of the mRNA level is

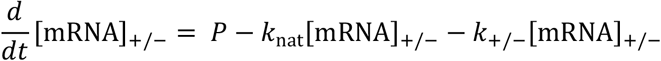

which results in a steady state mRNA concentration

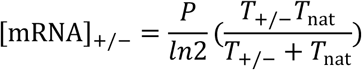

The dynamic range will therefore be

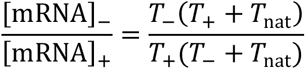

For gRNA activation, active gRNAs are produced from transcripts described by the equations shown above:

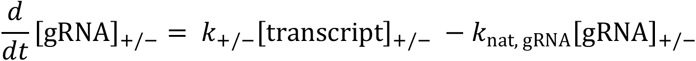

The transcript level with and without ligand is given by the formula derived above for the mRNA level. For the steady state, this simplifies to

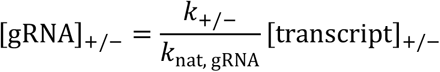

which yields a dynamic range of

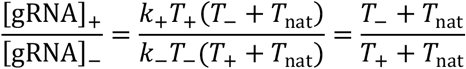

Depending on the half-lives for the various RNA species involved, this results in very different dynamic ranges (or on/off ratios) and maximum expression levels/activities. For instance, using half-lives of *T*_nat_ = 8h, *T*_−_ = 16*h* and *T*_+_ = 0.2*h* (8 hours is a typical half-life for a mammalian mRNA^34^) gives a dynamic range of 27.3 for mRNAs and 2.93 for gRNAs using this model. Using half-lives of *T*_nat_ = 1*h*, *T*_−_ = 32*h* and *T*_+_ = 0.4*h* (the stability of transcripts containing a Cas12a gRNA handle is likely much lower than that of a typical mRNA because the 5’ cap is removed, see also Supplementary Text 4) gives a dynamic range of 3.39 for mRNAs and 23.6 for gRNAs.

Next to the ON/OFF ratio, an important parameter is the maximum activity in the “ON” state – a good ON/OFF ratio is not useful if the switching occurs between very low overall expression levels. For mRNAs, this absolute performance can be quantified via the ratio of mRNA for the non-induced ribozyme and mRNA for the non-switchable construct, i.e.

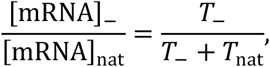

which becomes 1 for a ribozyme without leak activity (k_-_ → 0).

For gRNA, by contrast, the absolute performance is governed by the ratio of gRNA for the induced ribozyme and gRNA for the case where all transcript is immediately converted to gRNA:

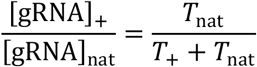

Plotting these values for different *T*_nat_, *T*_−_ = 16*h* and *T*_+_ = 0.4*h* yields

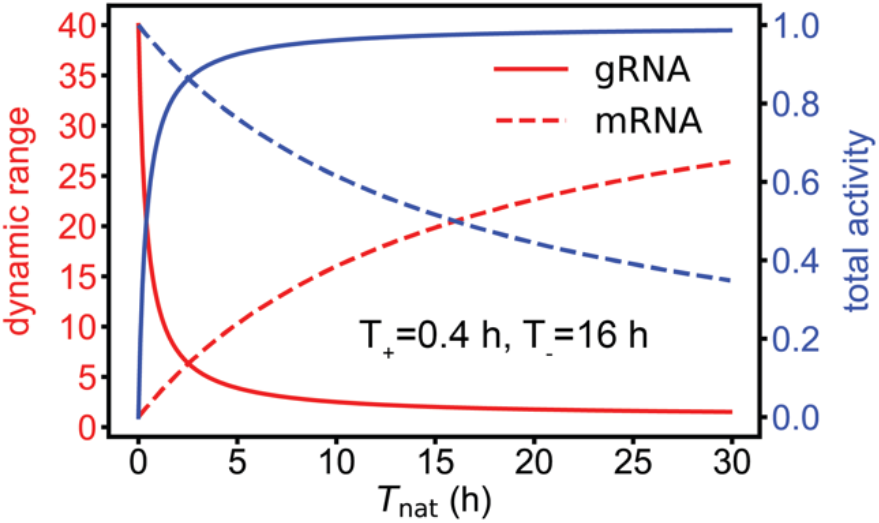

In both cases there is a trade-off between dynamic range and total activity that can be maintained. Even though this simple model is rather simplistic, it qualitatively illustrates the following point: The dynamic range of ribozyme-switchable mRNAs is highly sensitive to non-cleaved transcript (i.e., undesired protein production in the presence of ligand), while the dynamic range of gRNAs is highly sensitive to leaky processing (i.e., undesired protein production in the absence of ligand).

## Supplementary Note 2 – NUPACK design considerations

### Design of in vitro pseudoknot-based 5’H SD gRNA

The design is specified in the file “1_SDgRNA_in-vitro_mCerulean-sensor.np” and shown schematically in Supplementary Figure 4a. The toehold length is set to be 14 nt, and the variable portion of the switch has a length of 12 nt, for a total switch length of 16 nt. The switch and toehold sequences are set to be complementary to the mCerulean mRNA. The last four nucleotides of the switch (AAUU), which are derived from the Cas12a handle and therefore fixed are not present on the trigger. In the complex between the trigger and the SD gRNA, the last four nucleotides detach without being displaced due to formation of the pseudoknot, obviating any need for fixed sequences in the trigger. The toehold of the SD gRNA is given extra weight because we assumed that its secondary structure is likely to be the rate limiting factor in the displacement. Long stretches of a single nucleotide and an excessive AU or GC content are forbidden. The trigger contains a hairpin at the 5’ and 3’ end and the SD gRNA contains a trigger at its 5’ end. These hairpins suppress repetitive transcription by T7 RNA polymerase (cf. the Supplementary Information of Oesinghaus et al. for details)^19^.

### Design of in vivo pseudoknot-based 3’H SD gRNAs

The target of the gRNA in the in vitro design discussed above is constrained to be complementary to the first few nucleotides of the handle sequence and the sequence of the input molecule. For our switching principle, this constraint is irrelevant because we only use this gRNA for transcript processing. The constrained target therefore belongs to the gRNA that is located at the 3’ end of the transcript, which is anyway inactive (cf. Supplementary Figure 1): The gRNA located at the 5’ end is what ultimately interests us.

### gRNA t2 – 3’H SD t2

The design of the first 3’H SD gRNA for a synthetic trigger is specified in “2_pCAG_t2_3HSD-t2.np” and shown schematically in Figure 3a. The toehold size was set to 20 nt to maximize the interaction between the SD gRNA and its trigger and the “variable” portion of the switch has a length of 12 nt. Because the switch is designed to interact with the gRNA spacer sequence (target t2), its “variable” portion is fixed by the spacer sequence for this design. 53 nt upstream and 50 nt downstream of the expected transcript are included to avoid extensive interactions with these sequences. The weight of these sequences is set to be very low because we do not care about their structure beyond their interaction with the newly designed sequences. The toehold is set to have a GC content between 35% and 70% and has a large weight because we assumed that its secondary structure is likely to be the rate limiting factor in the displacement. The trigger includes the U6+27 sequence to increase its stability.

“3_pCAG_t2_3HSD-t2_noToe.np” designs a trigger with an identical switch but a non-complementary toehold.

The kinetic insulator design (“4_pCAG_t2_3HSD-t2_kinIns.np “) is virtually identical to the regular design except for two random nucleotides in the switch sequence. It is shown schematically in Supplementary Figure 5b. Rather than having a continuous switch stem of 16 nt (including the fixed AAUU sequence), its switch is divided into two times 7 nt with a 2 nt bulge in between.

### gRNA t1 - 3’H SD mScar

For these designs, the switch pairs with a 10 nt extension at the 3’ end of the gRNA spacer sequence t1 to avoid the sequence constraints of the design outlined above. The design is shown schematically in Supplementary Figure 6a. Two sequences were chosen by manual inspection of the 3’ UTR of the mScarlet mRNA (cf. Supplementary Figure 6b). The sequences were chosen based on 1) their ability to form structure-free triggers and toeholds, 2) having no more than three consecutive Cs or Gs to avoid G-quadruplexes, 3) their accessibility within the 3’ UTR, 4) their ending on AAT to displace a few of the fixed nucleotides in the switch (as shown in Figure 5a, this requirement is not strictly necessary). The toehold length was chosen to be 18 nt and the variable switch length was chosen to be 10 nt. The reduced toehold and switch length are due to a decreased chance of a structure-free trigger for longer natural sequences.

### gRNA t1 – 3’H SD mScar clamp and clamp-v2

For the clamp-v1 design, two times five nt complementary to the target sequence separated by a bulge were chosen by hand and placed at the 5’ end of the gRNA to reduce target-toehold interactions (cf. Supplementary Figure 6c). For the clamp-v2 design, a single 10 nt clamp was chosen. The clamp is separated from the handle by a 6 nt sequence designed by NUPACK to not interact with the rest of the SD gRNA (“5_pCAG_t1-3HSD-mScar-clamp-v2”). Additionally, 14 nt clamps at the 5’ end and 3’end of the SD gRNA pair the expected 5’ end and close to the expected 3’ end of the transcript. As seen in Supplementary Figure 8, outside of the sequence context of the transcript, both SD gRNAs have a structure-free toehold. Within the sequence context, the toehold of the clamp-v1 construct is almost fully paired while the clamp-v2 construct retains a structure-free toehold.

### gRNA t2 – 3’H SD seq1 to seq4

The design is specified in the files “6_pCAG_t2_3HSD-seq1.np” and “7_pCAG_t2_3HSD-seq2to4”. It is shown schematically in Figure 3d. 148 nt of the expected transcript sequence at the 5’ end and 154 nt of the expected transcript sequence at the 3’ end are included to optimize against interaction of the toehold or switch with these sequences. The t2 spacer sequence was 20 nt long and connected to the switch stem by a 3 nt separator. The switch has a length of 14 nt (including 4 fixed bases) and the toehold has a length of 20 nt. Stretches of more than three repeats of the same base were excluded from the toehold and switch sequences.

For the seq1 design, the toehold was set to have between 30% and 60% GC content. The switch did not have a fixed GC content and was allowed to contain wobble base pairs. For the seq2, seq3, and seq4 designs, the toehold was set to have between 25% and 60% GC content. The variable portion of the switch was set to have between 30% and 60% GC content and was not allowed to contain wobble base pairs. Again, both designs assign a large weight to the toehold because association between the toehold and trigger was assumed to be the rate-limiting factor for the displacement process.

## Supplementary Note 3 – Kinetic insulators and the mScarlet 3’ UTR as an input for 3’H SD gRNAs

In our previous SD gRNA design in bacterial cells, including a “kinetic toehold” in the switch sequence, i.e. leaving a few nucleotides of the switch unpaired, greatly increased the on/off ratio. We attempted the same for this design (Supplementary Figure 5b). Here, introduction of kinetic toeholds does not improve performance. Because we expect this design change to improve the speed of the strand displacement process, we suspect that the performance in this case might be limited by the initial association of the strands via the toehold.

We tested the unconstrained design without kinetic toeholds described in Figure 3 for two natural sequences derived from the 3’ untranslated region (UTR) of an mScarlet mRNA transcribed using the U6+27 promoter (Supplementary Figure 6a and b). One sequence failed to activate the 3’H SD gRNA, while the other activated it approximately 3-fold (Supplementary Figure 6a). Even for this sequence, activation by the full mScarlet mRNA was not significant.

A lack of toehold accessibility could be a possible reason for the failure of the first sequence because the target sequence interacts with the toehold sequence (Supplementary Figure 7). We tried to suppress this interaction with a clamp sequence (Supplementary Figure 6c). This improved activation by the U6-transcribed trigger to approximately 2-fold, which is still much less than desired. Again, lack of toehold accessibility in the full transcript might be responsible (Supplementary Figure 8). We introduced additional clamps to insulate the folding of the gRNA from the folding of the rest of the transcript (Supplementary Figure 6c, Supplementary Figure 8). This approach did not lead to further improvement of activation by the trigger.

## Supplementary Note 4 – The switching mechanism

We assume that various factors contribute to the switching mechanism of our conditional Cas12a guide RNAs, which depend on different cellular localization and degradation mechanisms. Our dCas12a-VPR is equipped with a nuclear localization factor and it is thus expected to be present both in the nucleus and in the cytoplasm, where it may bind to gRNAs with an accessible handle.

Because the 5’ end of the transcript is removed by dCas12a constitutively, either the large amount of sequence on the 3’ end could be responsible for the inhibition or, more specifically, the poly(A) tail might be necessary for the principle to work. In Figures S11a and b, we tested the influence of a hairpin (~23 nt), a long random sequence (~30 nt), or a combination of both on the 3’ end of the gRNA after processing for U6- and CAG-transcribed gRNAs. These extended gRNAs are inactive for CAG promoter transcripts, but not for U6 promoter transcripts. For the CAG-transcribed gRNAs, the construct with the random 30 nt sequence has less activity than the construct with the 23 nt hairpin. As shown in the preceding section, a 10 nt 3’ extension does not reduce the activity of gRNAs (cf. controls in Supplementary Figure 6a). RISC-based cleavage also failed to activate a construct retaining a hairpin. Since the U6- and CAG-based gRNAs should nominally be almost the same after processing, we believe that steric hindrance is unlikely to be the reason for the inactivity of constructs with a long 3’ extension.

Therefore, the history of the gRNAs prior to processing could be responsible for the difference in behavior. As expected, U6-transcribed gRNAs are mostly localized to discrete loci in the nucleus, while CAG-transcribed gRNAs are mostly localized to the cytoplasm (Supplementary Figure 10)^30^. Removal of the poly(A) tail could expose CAG-transcribed gRNAs to cytoplasmic nucleases, possibly leading to strong degradation. We tested this hypothesis using RT-qPCR (Supplementary Figure 11c), where we quantified the relative amount of gRNA with and without the presence of Cas12a. There is a large difference in the behavior of U6-transcribed and CAG-transcribed gRNAs upon addition of a Cas12a plasmid. For both U6 gRNAs, there is a large ~7-8 fold increase in the amount of RNA when Cas12a is present. For CAG gRNAs, there is a 5 to 10 fold reduction in the amount of RNA. This behavior could at least partially be explained by differences in intrinsic stability. We expect the U6 gRNAs transcripts to have a half-life of less than 30 minutes ^35,36^, while the CAG gRNAs should have a half-life of several hours^34^. For the U6 gRNAs, Cas12a therefore stabilizes the gRNA, while for CAG gRNAs, removal of the 5’ cap and poly(A) tail might make the transcript temporarily more vulnerable to degradation.

The difference in reduction of the amount of total gRNA for the different type of CAG constructs does not explain the observed differences in activity. However, since we only quantified the gRNA itself regardless of its sequence context, we do not know if processing actually occurs as expected for all constructs. To gain a more detailed insight into the state of the gRNAs, we performed RT-qPCR for two additional primers sets to also quantify the amount of total transcript and the amount of transcript that retains its 3’ tail (Supplementary Figure 11d). Notably, the total amount of gRNA in the absence of Cas12a is consistently shown by all three primer pairs to be lower for the active construct than for the inactive constructs, for unknown reasons. This difference is far too small to account for the measured difference in activity, however.

The standard constructs with and without a 3’ handle (3’H and no 3’H) show the expected behavior. For the 3’H construct, only a small amount of non-processed transcript remains when Cas12a is present. For the no 3’H construct, the transcripts have mostly been cleaved at the 5’ end, but retain their 3’ tail. The construct containing the 30 nt random sequence has approximately the same amount of completely unprocessed transcript as the regular 3’H construct, but much less fully processed gRNA. This could imply that the random sequence increases the vulnerability to degradation after processing, and possibly even after Cas12a is already bound, thereby explaining the reduced activity. The results for the construct containing the hairpin are more mixed. The gRNA appears to be slightly more vulnerable, but less so than for the long random sequence.

As a rough estimate for the amount of fully processed gRNA, we subtracted the amount of unprocessed or 3’ tail-containing transcript from the total gRNA amount (Supplementary Figure 11f). For the regular 3’H sample, the amount of gRNA is relatively high for both types of subtraction, indicating a large amount of fully processed gRNA. For the sample without a 3’ handle, the amount is high when the amount of unprocessed transcript is subtracted, but low when the amount transcript containing a 3’ tail is subtracted, indicating that transcripts have been processed at the 5’ end, but not at the 3’ end, as expected. For the samples which retain a long 3’ sequence after processing, the amount of remaining gRNA is low for both types of subtraction, implying that there is only very little fully processed gRNA.

Taken together, these results indicate that several mechanisms might be jointly responsible for switching depending on the precise design of the switched gRNA. For constructs without any 3’ tail cleavage (i.e. no 3’H), either continued localization in the cytoplasm or steric hindrance, but not degradation, are responsible for the absence of activity. For the constructs retaining a 20-30 nt long 3’ sequence, increased degradation seems likely to dominate the switching principle. Notably, these results indicate that the poly(A) tail is not strictly necessary for the switching principle, allowing to place multiple switchable gRNAs onto a single transcript.

## Notes

### Competing Interest Statement

The authors have declared no competing interest.

